# MorphoFeatures: unsupervised exploration of cell types, tissues and organs in volume electron microscopy

**DOI:** 10.1101/2022.05.07.490949

**Authors:** Valentyna Zinchenko, Johannes Hugger, Virginie Uhlmann, Detlev Arendt, Anna Kreshuk

## Abstract

Electron microscopy (EM) provides a uniquely detailed view of cellular morphology, including organelles and fine subcellular ultrastructure. While the acquisition and (semi-)automatic segmentation of multicellular EM volumes is now becoming routine, large-scale analysis remains severely limited by the lack of generally applicable pipelines for automatic extraction of comprehensive morphological descriptors. Here, we present a novel unsupervised method for learning cellular morphology features directly from 3D EM data: a convolutional neural network delivers a representation of cells by shape and ultrastructure. Applied to the full volume of an entire three-segmented worm of the annelid *Platynereis dumerilii*, it yields a visually consistent grouping of cells supported by specific gene expression profiles. Integration of features across spatial neighbours can retrieve tissues and organs, revealing, for example, a detailed organization of the animal foregut. We envision that the unbiased nature of the proposed morphological descriptors will enable rapid exploration of very different biological questions in large EM volumes, greatly increasing the impact of these invaluable, but costly resources.

## Introduction

Development of multicellular organisms progressively gives rise to a variety of cell types, with differential gene expression resulting in different cellular structures that enable diverse cellular functions. Structures and functions together represent the phenotype of an organism. The characterization of cell types is important to understand how organisms are built and function in general (***Arendt (2008)***). Recent advances in single-cell sequencing have allowed the monitoring of differential gene expression in the cell types that make up the tissues and organs of entire multicellular organisms, and led to the recognition of distinct regulatory programs driving the expression of unique sets of cell type-specific effector genes (***Hobert and Kratsios (2019)***; ***Sebé-Pedrós et al. (2018b,a)***; ***Musser et al. (2021)***; ***Fincher et al. (2018)***; ***Siebert et al. (2019)***). These distinct genetic programs make the identity and thus define cell types (***Arendt et al. (2016)***; ***Tanay and Sebé-Pedrós (2021)***). The genetic individuation of cell types then translates into phenotypic differences between cells, and one of the most intriguing questions is to determine this causation for each cell type. Only then will we understand their structural and functional peculiarities and intricacies and learn how evolutionary cell type diversification has sustained organismal morphology and physiology. As a prerequisite towards this aim, we need a comprehensive monitoring and comparison of cellular phenotypes across the entire multicellular body, which can then be linked to the genetically defined cell types.

Recent efforts have focused on large-scale high-resolution imaging to capture the morphological uniqueness of cell types. For the first time, the rapid progress in volume EM allows this for increasingly large samples, including organs and entire smaller animals, with nanometer resolution (***Bae et al. (2021)***; ***Vergara et al. (2021)***; ***Zheng et al. (2018)***; ***Scheffer et al. (2020)***; ***Cook et al. (2019)***; ***Verasztó et al. (2020)***). Supported by automated cell segmentation pipelines (***Heinrich et al. (2021)***; ***Macrina et al. (2021)***; ***Pape et al. (2017)***; ***Müller et al. (2021)***), such datasets allow characterization and comparison of cellular morphologies, including membrane-bound and membraneless organelles and inclusions, at an unprecedented level of detail for thousands of cells simultaneously (***Turner et al. (2022)***; ***Vergara et al. (2021)***). A notable challenge of such large-scale studies is the analysis of massive amounts of imaging data that can no longer be inspected manually. The latest tools for visual exploration of multi-terabyte 3D data allow for seamless browsing through the regions of interest (***Pietzsch et al. (2015)***; ***Maitin-Shepard et al. (2021)***), given such regions are spec-ified and a researcher has a clear study question in mind. However, if the goal is to characterise and/or compare morphologies of most or all cells in the dataset, one has to be able to automatically assign each cell an unbiased comprehensive morphological description - a feature vector with values representing the morphological properties of the cell in the most parsimonious manner.

The need for automation and quantitative evaluation has triggered the design of complex engineered features that capture specific cell characteristics (***Barad et al. (2022)***). However, such features often have to be manually chosen and mostly provide only a limited, targeted view of the overall morphology. To alleviate this problem, intricate modelling pipelines for cellular morphology quantification have been developed (***Ruan et al. (2020)***; ***Driscoll et al. (2019)***; ***Phillip et al. (2021)***), still these are often data- and task-specific. For unbiased monitoring and comparison of cellular morphologies, broadly applicable automated pipelines are missing. They should in addition be capable of extracting comprehensive representations that are not biased by manual feature selection, the end task or the type of data used. We expect that developing such techniques will not only speed up the process of scientific discovery in big volume imaging datasets, but also facilitate detecting unusual and underrepresented morphologies.

In the last decade artificial neural networks have been shown to exceed manually engineered pipelines in extracting comprehensive descriptors for various types of data, and to be particularly suited for the extraction of visual features (***Krizhevsky et al. (2012)***; ***Falk et al. (2019)***; ***Simonyan and Zisserman (2014)***; ***Girshick et al. (2014)***). Moreover, the rise of so-called self-supervised training methods has removed the necessity for generating massive amounts of manual annotations previously needed to train a network. Besides the obvious advantage of being less labour-intensive, self-supervised methods do not optimise features for a specific end-task, but instead produce descriptors that have been shown to be useful in a variety of downstream tasks, such as image classification, segmentation and object detection (***He et al. (2020)***; ***Van den Oord et al. (2018)***; ***Tian et al. (2020)***). Such methods have already shown to be successful for phenotyping cells in image-based profiling (***Lu et al. (2019)***; ***Lafarge et al. (2019)***) and for describing local morphology of patches or subcompartments within neural cells in 3D electron microscopy of brain volumes (***Huang et al. (2020)***; ***Schubert et al. (2019)***). Building on the latest achievements, we aim to expand the feature extraction methods to automated characterisation of cellular morphology at the whole animal level.

Here we present the first framework for the fully unsupervised characterization of cellular shapes and ultrastructures in a whole-body dataset for an entire animal. We apply this new tool to the fully segmented serial blockface electron microscopy (SBEM) volume of the 6 days post fertilisation young worm of the nereid Platynereis dumerilii, comprising 11,402 mostly differentiated cells with distinct morphological properties (***Vergara et al. (2021)***). We show that our method yields morphological descriptors - *MorphoFeatures* - that are in strong agreement with human perception of morphological similarity and quantitatively outperform manually defined features on cell classification and symmetric partner detection tasks. We further illustrate how our features can facilitate the detection of cell types by morphological means, via similarity-based clustering in the MorphoFeature vector space, sorting the cells into morphologically meaningful groups that show high correlation with genetically defined types such as muscle cells and neurons. Our pipeline also allows for the characterization of rare cell types such as enteric neurons and rhabdomeric photoreceptors, and detects distinct cell populations within the developing midgut. Finally, defining feature vectors that also represent the morphology of immediate neighbours, we group cells into tissues, and also obtain larger groupings that represent entire organs. We show that such neighbourhood-based *MorphoContextFeature* vector clustering reproduces manually annotated ganglionic nuclei in the annelid brain, and represents a powerful tool to automatically and comprehensively detect the distinct tissues that belong to and make up the foregut of the nereid worm - including highly specialised and intricate structures such as the neurosecretory infracerebral gland (***Baskin (1974)***; ***Hofmann (1976)***; ***Golding (1970)***) and the axochord that has been likened to the chordate notochord (***Lauri et al. (2014)***). We further show that such morphologically defined tissues and organs correlate with cell type and tissue-specific gene expression. Our work thus sets the stage for linking genetic identity and structure-function of cell types, tissues and organs across an entire animal.

The MorphoFeatures for the Platynereis dataset, as well as the code to generate and analyse them are available at https://github.com/kreshuklab/MorphoFeatures.git.

## Results

### Unsupervised deep learning extracts extensive morphological features

Our pipeline has been designed to extract morphological descriptors of cells (MorphoFeatures) from EM data. It requires prior segmentation of all the cells and nuclei of interest. The pipeline utilises segmentation masks to extract single cells/nuclei and represents their morphology as a combination of three essential components: shape, coarse and fine texture (Figure 1A). We train neural networks to represent these components independently, using point clouds as input for shape, low resolution raw data for coarse texture with sufficient context and small, high resolution crops of raw data for fine texture. Separating these components ensures they are all represented in the final features, and facilitates further interpretation. The features are combined to form one feature vector for each cell of the animal.

**Figure 1.**
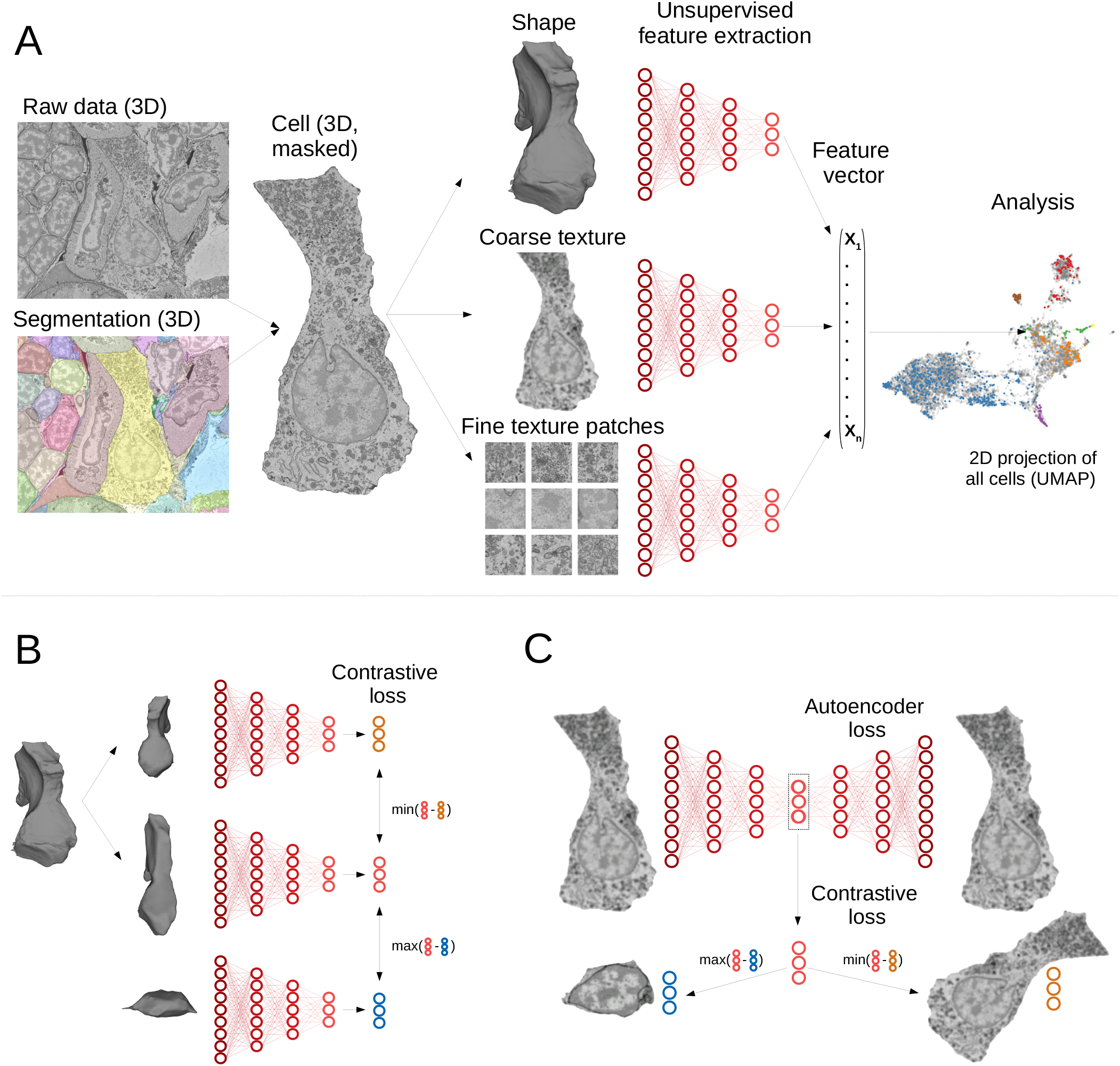
Deep learning pipeline for extracting MorphoFeatures. **A**. Cell segmentation is used to mask the volume of a specific cell (and its nucleus) in the raw data. Neural networks are trained to represent shape, coarse and fine texture from the cell volume (separately for cytoplasm and nuclei). The resulting features are combined in one MorphoFeatures vector that is used for the subsequent analysis. **B**. Training procedure for the shape features. A contrastive loss is used to decrease the distance between the feature vectors of two augmented views of the same cell, and increase the distance to another augmented cell. **C**. Training procedure for the texture features. Besides the contrastive loss, an autoencoder loss is used that drives the network to reconstruct the original cell from the feature vector

A common way to train a neural network for extracting relevant features is to use some prior information about samples in the dataset as a source of supervision. For example, to extract morphological representations of cells one could use a complementary task of predicting cell lines (***Yao et al. (2019)***; ***Doan et al. (2020)***; ***Eulenberg et al. (2017)***), experimental conditions the cells are coming from (***Caicedo et al. (2018)***), or classification of proteins fluorescently labelled in the cells (***Kobayashi et al. (2021)***). However, such metadata is mostly unavailable for EM volumes, and manual annotation might become infeasible with increasing data size. Moreover, exploratory studies are often aimed at discovering unusual morphology, new cell types or tissues. In this case, defining supervision is not only difficult, but might also bias the exploration towards the “proxy” groups used for supervision. To enable immediate data exploration even for datasets where direct supervision is hard to define and the ground truth cannot easily be obtained, we developed a fully unsupervised training pipeline that is based on two complementary objectives (Figure 1B,C). The first is an autoencoder reconstruction loss, where a network extracts a low-dimensional representation of each cell and then uses this representation to reconstruct back the original cell volume. This loss encourages the network to extract the most comprehensive description. The second objective is a contrastive loss that ensures the feature vectors extracted from two similarly looking cells (positive samples) are closer to each other than to feature vectors extracted from more dissimilar cells (negative samples). Since we do not know in advance which samples can be considered positive for our dataset, we are using slightly different views of the same cell, generated by applying realistic transformations to cell volumes (see Methods). The combination of these two losses encourages the learned features to retain the maximal amount of information about the cells, while enforcing distances between feature vectors to reflect morphological similarity of cells.

The pipeline was trained and applied on the cellular atlas of the marine annelid Platynereis dumerilii (***Vergara et al. (2021)***). It comprises a 3D serial block-face electron microscopy volume of the whole animal that has sufficient resolution to distinguish ultrastructural elements (organelles and inclusions, nuclear and cytoplasm texture, etc.) and an automated segmentation of 11,402 cells and nuclei (Figure 1A). Additionally, whole-animal gene expression maps are available that cover many differentiation genes and transcription factors.

Such a high number of morphologically and genetically diverse cells can make the initial exploratory analysis difficult. We designed MorphoFeatures to enable unbiased exploration of morphological variability in the animal on both cellular and tissue level; and indeed our training procedure yields 480 features that extensively describe each cell in terms of its cytoplasm and nucleus shape, coarse and fine texture (80 features for each category). The features can be found at https://github.com/kreshuklab/MorphoFeatures.git, the feature table can be directly integrated with the Platynereis atlas of ***Vergara et al. (2021)*** through the MoBIE plugin in Fiji (***Schindelin et al. (2012)***) (https://zenodo.org/record/2602755).

### MorphoFeatures allow for accurate morphological class prediction

Good morphological features should distinguish visibly separate cell groups present in the data. To estimate the representation quality of our MorphoFeatures we quantified how well they can be used to tell such groups apart. We took the morphological cell class labels available in the dataset, proofread, corrected and expanded them. These labels include 7 general classes of cells: neurons, epithelial cells, midgut cells, muscle cells, secretory cells, dark neurosecretory cells and ciliary band cells (Figure 2A) with ∼60 annotated cells per class in the whole volume. It is worth noting that these cell classes were chosen based on visual morphological traits and do not necessarily represent genetically defined cell type families (***Arendt et al. (2019)***). For example, while ciliary band cells are a small narrow group of morphologically homogeneous cells that likely constitute one cell type family, secretory cells were defined as cells with abundant organelles/inclusions in their cytoplasm that could indicate secretion, spanning a wider range of morphologies and, potentially, genetically-defined cell types.

**Figure 2.**
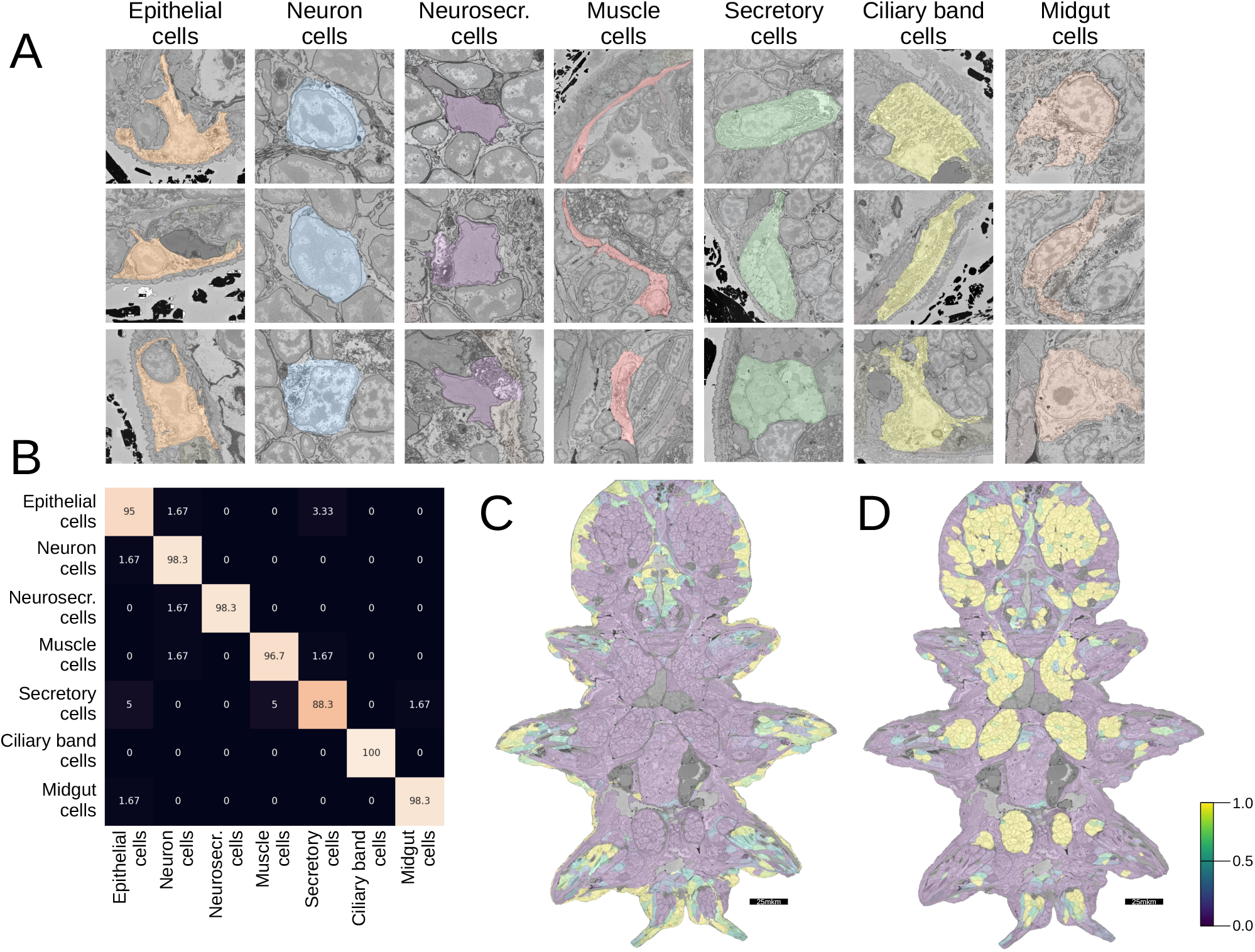
Morphological class prediction. **A**. Examples of cells from 7 manually defined morphological classes used for evaluation. **B**. Confusion matrix of class prediction for the logistic regression model. Rows are the labels, columns are the predictions. **C**. The predicted probability of the epithelial class in the whole animal. Note, that while a few cells on the animal surface have been used for training of the logistic regression, no labels were given in the gut opening or chaetae which are still correctly recognized as epithelial. **D**. The predicted probability of neural cells in the whole animal.

We extracted MorphoFeatures by passing all the cells through the neural network trained as described above (Figure 1A) and used them as an input to train a logistic regression classifier to predict the above-mentioned cell classes. This classifier has few parameters and can be trained even on the modest amount of labels we have available. If any subset of the features strongly correlates with a given cell class, the model will show high performance. A classifier with MorphoFeatures achieved an accuracy of 96%, showing that the learned features are sufficient to distinguish broad morphological classes. To better understand remaining inaccuracies, we show the errors made by the classifier in Figure 2B and Figure S1. This reveals, as expected, that while morphologically homogeneous classes can be easily separated, the classifier tends to confuse some secretory cells with other types (midgut and epithelial cells) that are also difficult to discern by eye and might well be mis-annotated. It also confuses the general neuron class with neurosecretory cells that represent a subclass of neurons.

To illustrate large-scale performance of the broad cell type classification model, we show its predictions on the whole animal in Figure 2C,D and Figure S1B. These results demonstrate that MorphoFeatures are sufficiently expressive to enable training a generalizable model from a small number of labels. Taking epithelial cells as a showcase: while the model was trained exclusively on the cells of the outer animal surface, it successfully predicted the cells outlining the gut opening and the chaetae cells as epithelial as well. Even more convincingly, the model predicted some cells in the midgut region as neurons, which after careful visual examination were indeed confirmed to be enteric neurons.

In ***Vergara et al. (2021)***, morphological classification of cells was performed based on manually defined features, including descriptors of shape (e.g., volume, sphericity, major and minor axes), intensity (e.g., intensity mean, median and standard deviation) and texture (Haralick features) extracted from the cell, nucleus and chromatin segmentations, with 140 features in total. To compare these explicitly defined features with implicitly learned MorphoFeatures, we passed them through the same evaluation pipeline yielding a classifier which achieves the accuracy of 94%, in comparison to 96% achieved by MorphoFeatures. The classifier mostly made similar mistakes, but performed worse on the classes of muscle and epithelial cells. Both sets of features demonstrate high accuracy on the broad morphological type classification task. For a more detailed comparison, we turn to a different task which allows us to implicitly, but quantitatively, evaluate how well the features group similar cells together. We use the metric introduced in ***Vergara et al. (2021)*** that is based on the fact that the animal is bilaterally symmetric, thus almost all the cells on one side of the animal will have a symmetric partner. The metric estimates how close a given cell is to its potential symmetric partner in the morphological feature space in comparison to all the other cells in the animal (see Methods). According to this metric our MorphoFeatures are 38% more precise in locating a symmetric partner than the ones from ***Vergara et al. (2021)***, showing the widely accepted superiority of neural network extracted features to explicitly defined ones.

### MorphoFeatures correspond to visually interpretable morphological properties

Neural networks are often referred to as “black boxes” to signify that it is not straightforward to trace learned features or decisions to specific input properties. To examine whether it is possible to understand which properties of cells were learned by our network, we looked at the groups of extracted MorphoFeatures (coarse and fine texture and shape features). For a set of features in each group we visualised cells that correspond to the maximum and minimum value of the corresponding feature (Figure 3). Visual inspection showed that many of them can be matched to visually comprehensible properties. For example, one of the cytoplasm coarse texture features (Figure 3, upper left) shows its minimal value in cells with a thin layer of cytoplasm surrounding the nucleus and its maximal value in wide muscle cells with prominent fibres and mitochondria. One of the nuclear coarse texture features (Figure 3, upper right) differentiates between big round nuclei with a rough surface and smaller extended ones. For fine texture, a cytoplasm-related feature (Figure 3, middle left) has its minimal value found in cells with medium-sized mitochondria and Golgi cisternae and its maximum correlated with small dark stretches of cytoplasm in dark neurosecretory cells. One of the nuclear fine texture features (Figure 3, middle right) is the lowest in cells with dark homogenous nuclei and has its peak in nuclei with a distinct dark nucleolus. For shape features of cellular membranes, we observed that some have one extreme in compact roundish surfaces such as neuron somata and the other extreme in long, straight and thin surfaces such as muscle cells (Figure 3, lower left). Shape features of nuclear membranes have some of their minima and maxima in compact deformed shapes and straight elongated or flat shapes (Figure 3, lower right).

**Figure 3.**
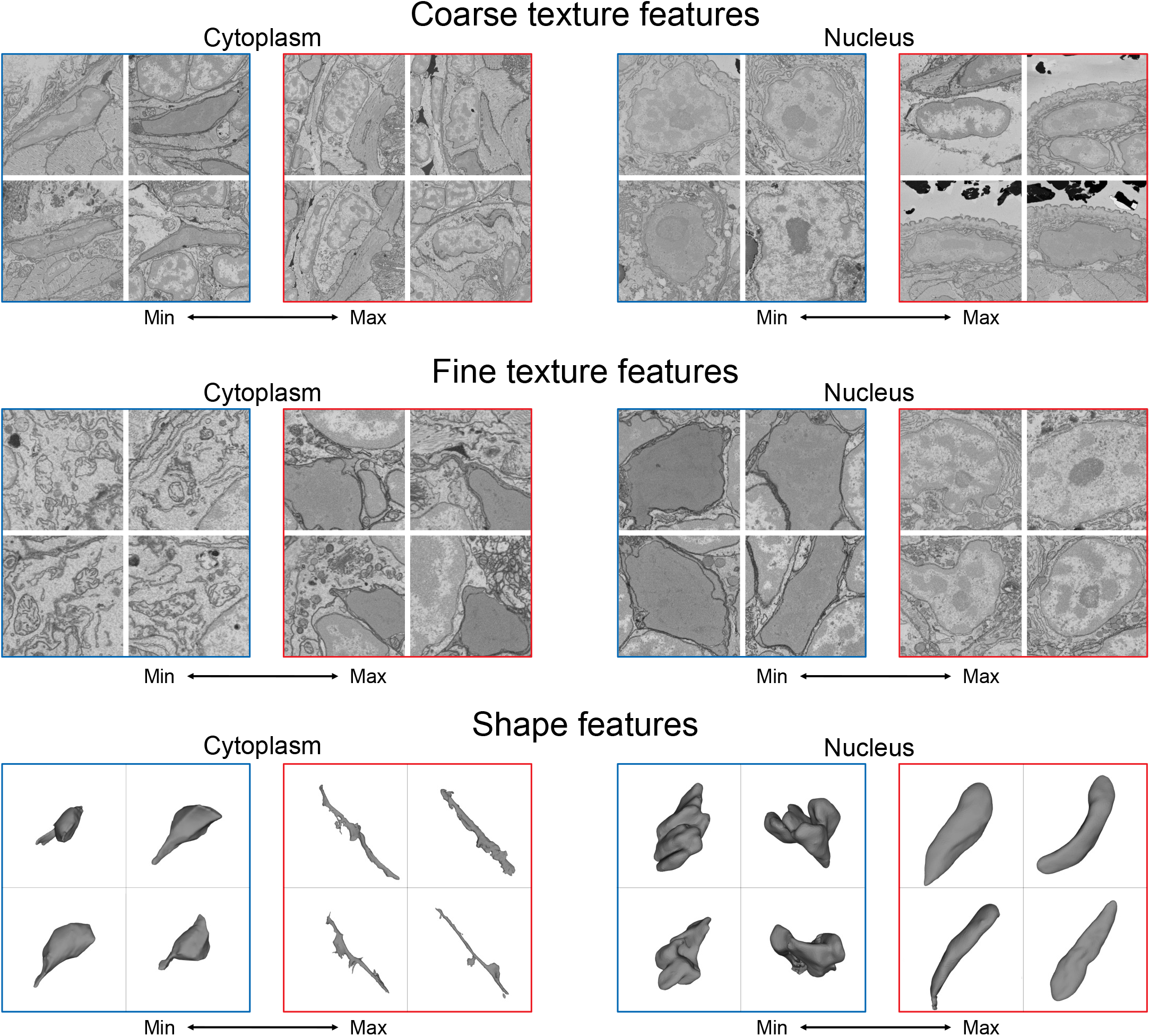
Visualization of the learned features. For each feature, four cells with a minimal (blue) and four cells with a maximal (red) value of the feature are shown, see text for detailed analysis.

### MorphoFeatures uncover groups of morphologically similar cells

Comprehensive analysis of morphological features should enable the grouping of cells with similar morphology, which may represent cell types. We investigated the grouping induced by MorphoFeatures by projecting the features of all the animal cells into 2D space using the dimensionality reduction technique UMAP (***McInnes et al. (2018)***). To explore the quality of our representations, we took a cell from the animal and visualised its 3 closest neighbours (Figure 4A). This indicates that cells in close proximity in the feature space appear highly visually similar and might represent cell types at the morphological level. For example a group of secretory cells in which the cytoplasm is filled with round electron-transparent vesicles (upper green panel) corresponds to the group of ‘bright droplets’ cells from ***Verasztó et al. (2020)***. Another group of secretory cells have an extensive ER surrounding the nucleus with a prominent nucleolus (lower green panel) and represents parapodial spinning gland cells (***Verasztó et al. (2020)***). One can also see clear groups of flat developing midgut cells (brown panel), short muscles (red panel), ciliary band cells on the outer animal surface (yellow panel) as well as extended flat and short round epithelial cells (orange panels). Another interesting observation is that even though the neurite morphology is not taken into account, neurons show remarkable diversity based on soma morphology only (blue panels). Careful examination of a group of cells, initially labelled as neurons, that are located next to epithelial cells on the UMAP projection (upper right blue panel), revealed that these cells are sensory cells that extend their processes to the epithelial surface.

**Figure 4.**
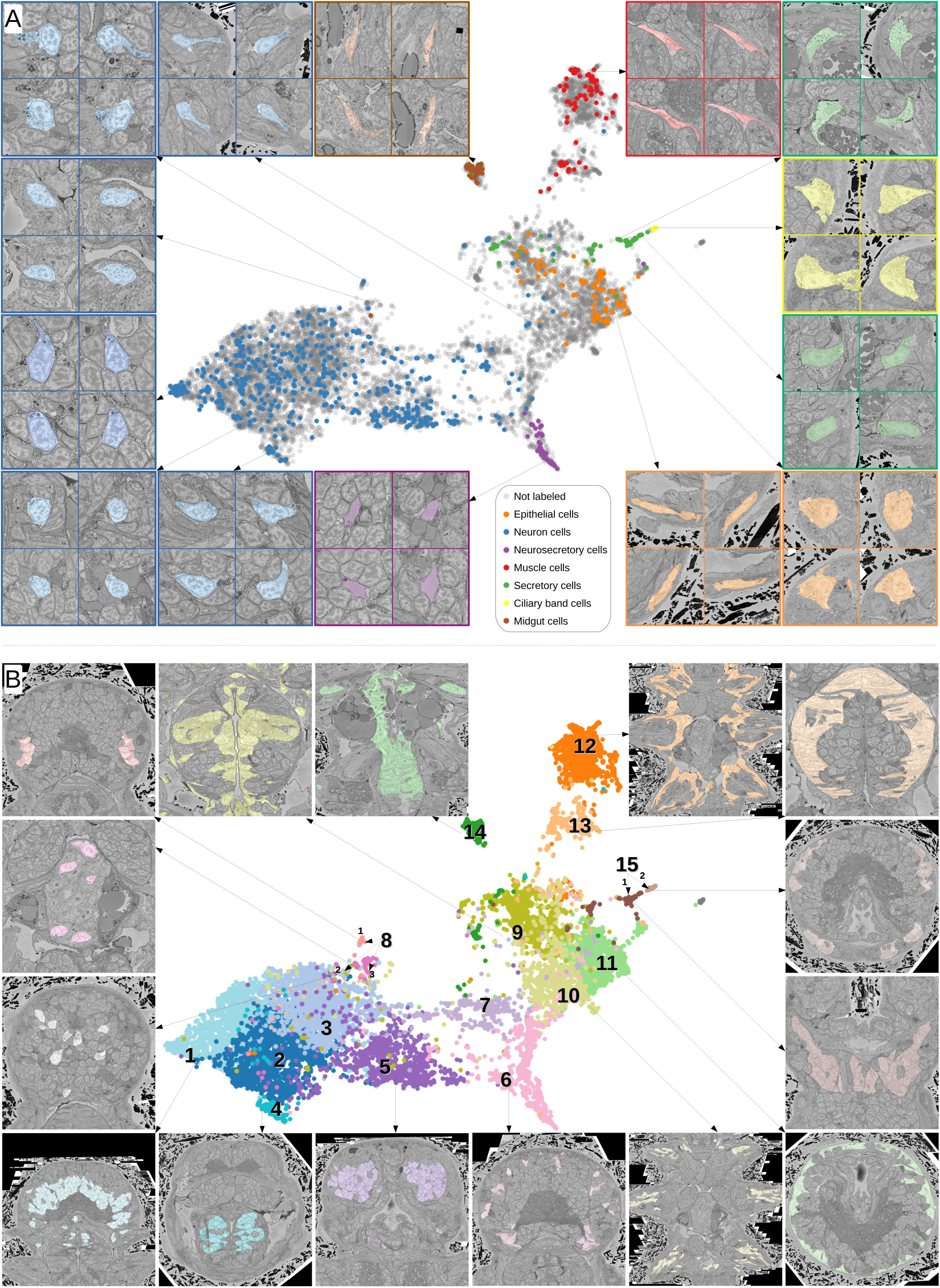
**A**. Finding visually similar cells using MorphoFeatures. Multidimentional features are visualized in 2D using UMAP. Each point represents a feature vector of a cell from the dataset. The cells for which annotations are available are visualized in respective colors. For a random cell, the cell and its three closest neighbors in the UMAP space are visualized in the EM volume. **B**. Visualising morphological clusters. Clustering results are visualised on the MorphoFeatures UMAP representation. For some clusters the cells comprising the cluster are shown in the animal volume to visualise the cell type. For example, cluster 6 precisely picks out the dark neurosecretory cells, while cluster 14 corresponds to the midgut cells (see Text for more details).

To explore the structure of the MorphoFeatures space, we performed hierarchical clustering of all cells. In the first clustering round we split cells into 15 broad morphological groups (Figure 4B). Cluster content could be revealed through available annotations and visualisation with the PlatyBrowser (***Vergara et al. (2021)***) (Figure 4B): Clusters 1-8 represent different groups of neurons; 9-11 epithelial cells; 12 and 13 were muscles; 14 midgut cells; while cluster 15 was further subclustered into secretory cells (subcluster 1) and ciliary band (subcluster 2). MorphoFeatures can thus guide the morphological exploration of a whole-animal volume, facilitating the discovery and annotation of morphologically coherent groups of cells (Figure 4B). For example, the foregut epithelial cells are distinct from the epithelial cells of the outer animal surface and chaetae cells (clusters 9, 11 and 10, respectively). Separate groups are also formed by different types of neurons, for example, foregut neurons (cluster 4).

### Morphological clusters have distinct gene expression profiles

To further describe the morphological clusters we took advantage of the whole-animal cellular gene expression atlas available for Platynereis (***Vergara et al. (2021)***), containing more than 200 genes mapped onto the EM volume. We looked for genes that are highly expressed in a given cluster, while also being specific for this cluster - having low expression in the other cells of the animal.

Many of our clusters show a clear genetic signature (Figure 5A, Figure S2). Among the neurons, cluster 5 shows the most specific gene expression including the Transient receptor potential (Trp) channels *pkd2, trpV4* and *trpV5*, and the bHLH transcription factor *asci*, which demarcate sensory cells of the palpal, antennal, and cirral ganglia (***Vergara et al. (2021)***). Cluster 1 shows specific expression of the homeodomain transcription factors *lhx6* and *phox2* and an *ap2* family member, while cluster 6 composed of dark neurosecretory cells shows specific expression of two neurosecretion markers, atrial natriuretic peptide receptor *anpra* and prohormone convertase *phc2*, identifying these as brain parts of the circular and apical nervous system, respectively (***Vergara et al. (2021)***; ***Arendt (2021)***). Cluster 8, another neural cluster, is enriched for the oxidative stress marker *cytoglobin* (*globin-like*) (***Song et al. (2020)***) and the bHLH transcription factor *mitf*. Subclustering revealed 3 subclusters, one of which represents rhabdomeric photoreceptors of the adult eye (Figure 4B, subcluster 8.1) that specifically express *mitf* and *globin-like*. An average cell in this subcluster (Figure 5B) shows abundant black inclusions. The second subcluster (8.2) contains pigment cells with black pigment granules. Cells in the third subcluster (8.3) are located in the midgut, with a smooth oval shape, neurite extensions and nuclei taking up the most cell volume (Figure 5C,D) - in striking contrast to other midgut cells but resembling neurons, making it likely that these cells represent enteric neurons previously postulated on the basis of serotonin staining (***Brunet et al. (2016)***).

**Figure 5.**
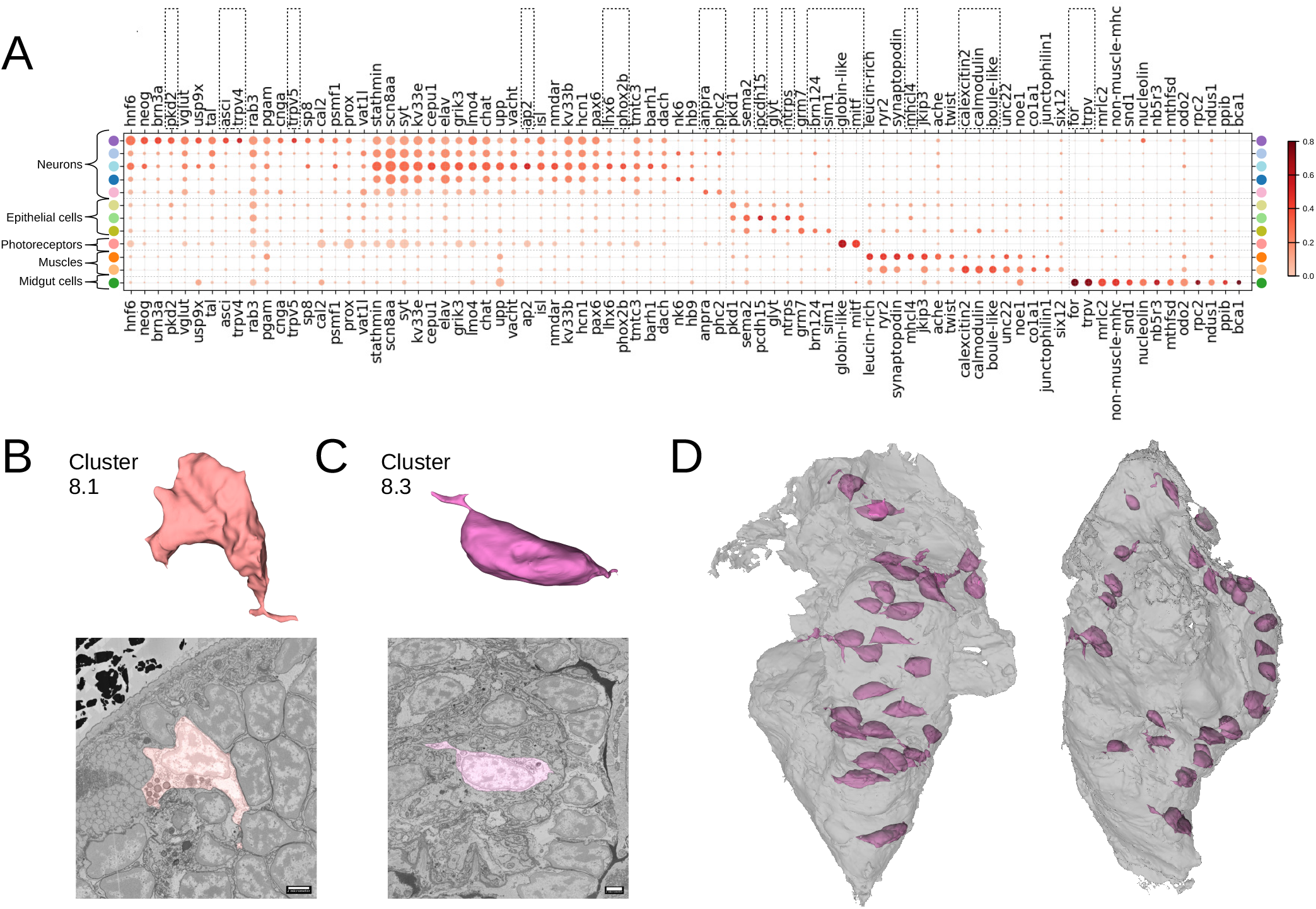
Clustering and gene analysis. **A**. Gene expression dot plot. The size of dots shows how much of the cluster expresses a gene; the colour shows how much of the expression of the gene is confined to a cluster (see Methods). The genes mentioned in the text are enboxed. The clusters lacking highly specific gene expression were not included. **B-C**. The average shape and texture (see Methods) rhabdomeric photoreceptors (cluster 8.1) and (C) the enteric neurons (cluster 8.3). **D**. Localisation of the enteric neurons (pink) in the midgut volume (grey). Left: frontal view, right: side view.

Among the epidermal cells, the outer cluster 11 is enriched for *protocadherin* (*pcdh15*) and *neurotrypsin* (*ntrps*), whereas cluster 9 representing foregut epidermis shows high specificity for the bHLH transcription factor *sim1* and the homeodomain factor *pou3* (*brn124*) (known to be expressed in stomodeal ectoderm in sea urchin (***Cole and Arnone (2009)***). For the muscles, the specific expression of striated muscle-specific myosin heavy chain *mhcl4* in cluster 12 denotes the group of striated somatic muscles, while its absence in cluster 13 indicates that the foregut muscles have not yet switched to expressing striated markers (***Brunet et al. (2016)***). Instead, the high and specific expression of the calexcitin-like sarcoplasmic calcium-binding protein *Scp2* (*calexcitin2*) (***White et al. (2011)***), *calmodulin* and of the Boule homolog *Boll* (*boule-like*) in cluster 13 suggests a calcium excitation and/or sequestration mechanism specific for foregut muscles. Finally, the cluster of midgut cells (cluster 14) specifically expresses the forkhead domain transcription factor *foxA* (*for*), a gut developmental marker (***Boyle and Seaver (2008)***) and *TrpV-c*, another Trp channel paralog of the vanilloid subtype with presumed mechanosensory function.

Beyond that, we noted considerable genetic heterogeneity in the midgut cluster. Subclustering (Figure 6) revealed one subcluster with strong expression of the smooth muscle markers *nonmuscle-mhc* and *mrlc2*, and another expressing a chitinase related to chitotriosidase (*nov2*) and the zinc finger transcription factor *Prdm16/mecom*, known to maintain homeostasis in intestinal epithelia (***Stine et al. (2019)***). Visualising the cells belonging to each subcluster and the specifically expressed genes in the PlatyBrowser (Figure 6E) revealed distinct territories in the differentiating midgut, which we interpret as midgut smooth musculature and digestive epithelia. We also detected an enigmatic third cluster located outside of the midgut, in the animal parapodia, comprising cells that resemble midgut cells morphologically (Figure 6D). For this subcluster, the current gene repertoire of the cellular expression atlas did not reveal any specifically expressed gene.

**Figure 6.**
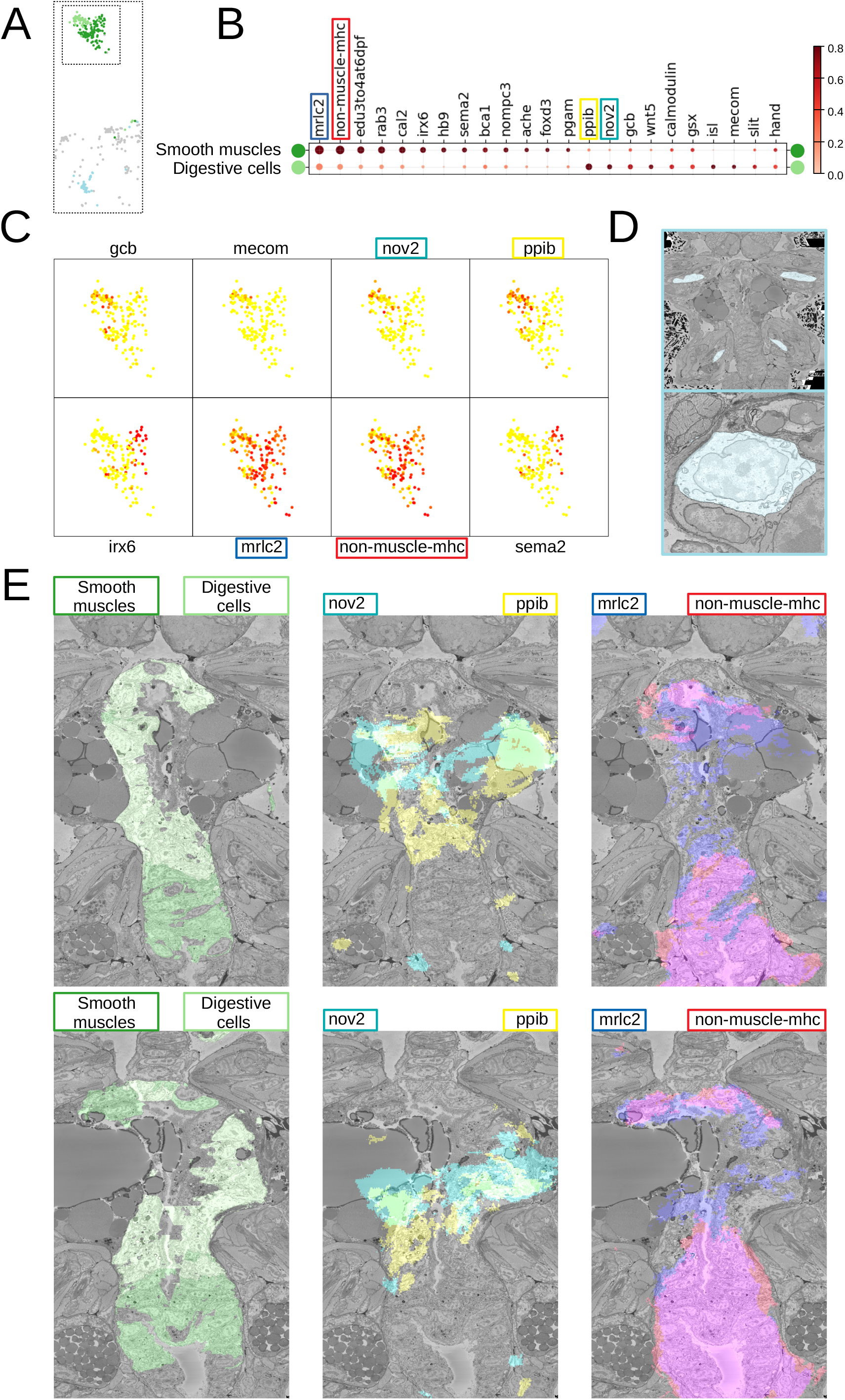
Midgut cell types with defining genes. **A**. Finer clustering of the midgut cluster results in three subclusters. **B**. Gene expression dot plot of the two midgut subclusters that are presumably developing smooth muscles and digestive cells of the midgut. The size of dots shows how much of the cluster expresses a gene; the colour shows how much of the expression of the gene is confined to a cluster (see Methods). **C**. Some of the genes shown to be differentially expressed in the two subclusters, plotted on the UMAP representation. **D**. The location (upper panel) and an example cell (lower panel) of the subcluster located in the animal parapodia. **E**. Cells belonging to the two cell types (left panels) and the genes differentiating them (center and right panels) are visualised in the animal volume, with color representing gene expression overlayed on the EM plane view.

### Adding neighbour morphological information helps to identify tissues and organs

Most cells do not function in isolation but rather form tissues, i.e. groups of structurally and functionally similar cells with interspersed extracellular matrix. We set out to systematically identify tissues with a new feature vector assigned to each cell that takes into account information about the morphology of neighbouring cells. For this, we combined the MorphoFeatures of a cell with the average MorphoFeatures of its immediate neighbours, yielding a feature vector that represents the morphology of both the cell and its surrounding, which we refer to as MorphoContextFeatures. Applying the representation analysis described before, we find that proximity in MorphoContextFeature space no longer reflects morphological similarity only, but rather identifies neighbouring groups of morphologically similar cells (whereby the neighbouring groups can be morphologically dissimilar). In other words, we now identify different tissues that together constitute organs composed of various tissues (Figure 7). For example, a separate assembly of groups of cells in the lower part of the UMAP projection represents the animal foregut subdivided into its constituting tissues comprising foregut neurons (lower left blue panel), foregut muscles (lower red panel) and foregut epithelium (left orange panel). Next to this we identify a group of muscles surrounding the foregut (middle left red panel).

**Figure 7.**
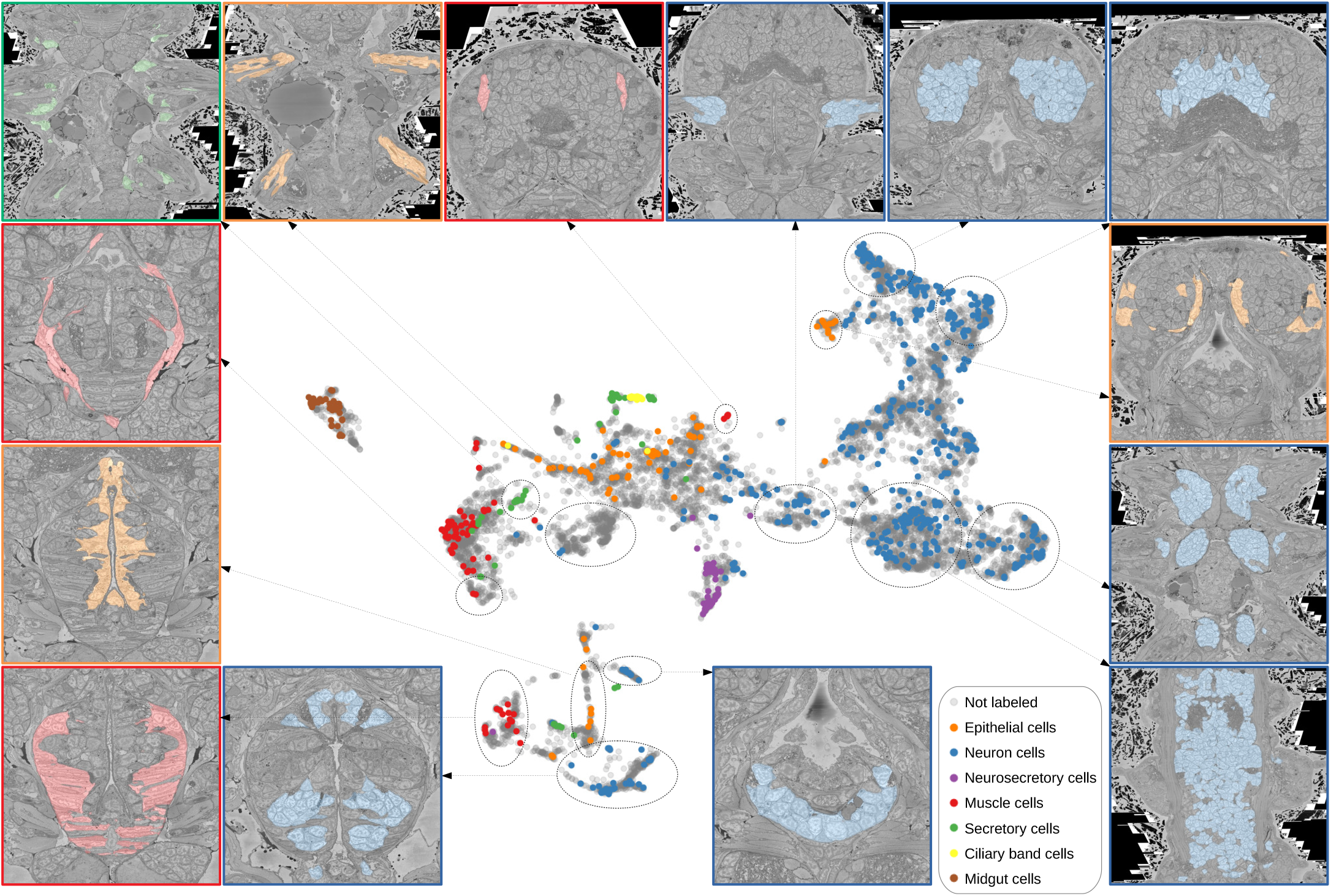
Characterizing neighbourhoods with MorphoContextFeatures. Panel colors represent classes cells belong to. Upper panels (from left to right): secretory (green) and epithelial (orange) cells of parapodia, antennal muscles that participate in moving and retracting antenna (red), cirral (blue), palpal (blue) and dorsal (blue) ganglia. Lower panels: foregut muscles (red), foregut neurons (blue), infracerebral gland (blue), ventral nerve cord (blue). Left panels: muscles surrounding the foregut (red), foregut epithelium (orange). Right panels: epithelial-sensory circumpalpal cells (orange) and peripheral ganglia (blue).

For the nervous system, adding neighbourhood information helps distinguishing cirral, palpal and dorsal ganglia (upper blue panels from left to right), as well as the ventral nerve cord and peripheral ganglia (lower and middle right blue panels). To benchmark the quality of this grouping, we clustered all the cells in the MorphoContextFeature space (see Methods), and compared the neuron clusters to both manually segmented ganglionic nuclei and to the genetically defined neuronal clusters taking into account available gene expression in the cellular atlas (***Vergara et al. (2021)***) (Figure 8A,B). Notably, manual segmentation of brain tissues relied on visible tissue boundaries, thus has lower quality in the areas where such boundaries are not sufficiently distinct.

**Figure 8.**
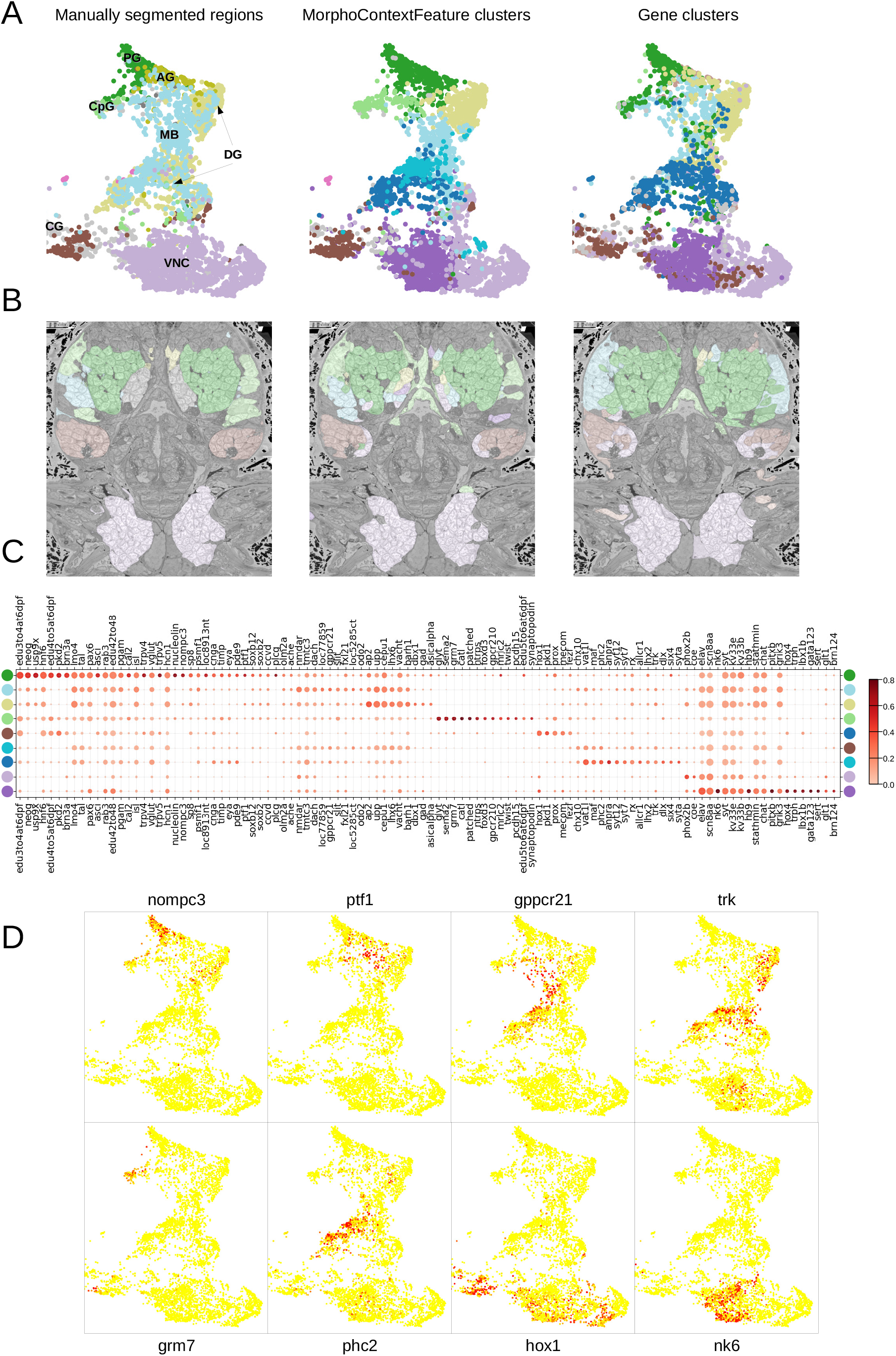
MorphoContextFeatures define ganglionic nuclei. **AB**. The animal ganglia as defined by manual segmentation (***Vergara et al. (2021)***), our MorphoContextFeature clustering and gene expression clustering displayed on (A) the UMAP representation and (B) in the animal volume. **C**. Gene expression dot plot of the ganglia defined by MorphoContextFeature clustering. The size of dots shows how much of the cluster expresses a gene; the colour shows how much of the expression of the gene is confined to a cluster (see Methods). **D**. Some of the genes shown to be differentially expressed in the ganglia defined by MorphoContextFeature clustering, plotted on the UMAP representation.

In essence, all three ways of defining ganglia lead to very similar results (Figure 8B), yet MorphoContextFeature clustering appeared to be the most powerful. Gene clustering failed to distinguish the circumpalpal ganglion, defined both by manual segmentation and MorphoContextFeature clustering, due to the lack of distinct gene expression in the atlas. Manual segmentation failed to distinguish between the dorso-posterior and dorsal-anterior ganglionic nuclei due to the lack of distinct tissue boundaries. These are well defined by gene clustering, and subdivided even further by MorphoContextFeature clustering.

### Unbiased exploration of foregut tissues

To further illustrate how MorphoContextFeatures can assist an unbiased exploration of an animal EM volume, we focused on the foregut. Subclustering whenever suitable (Figure 9A) (see Methods), the resulting clusters revealed various foregut tissues including foregut epidermis, ganglia, inner musculature and a prominent muscle surrounding the foregut. We also found a tissue of secretory neuron-like cells that surround the foregut like a collar (Figure 9B,C).

**Figure 9.**
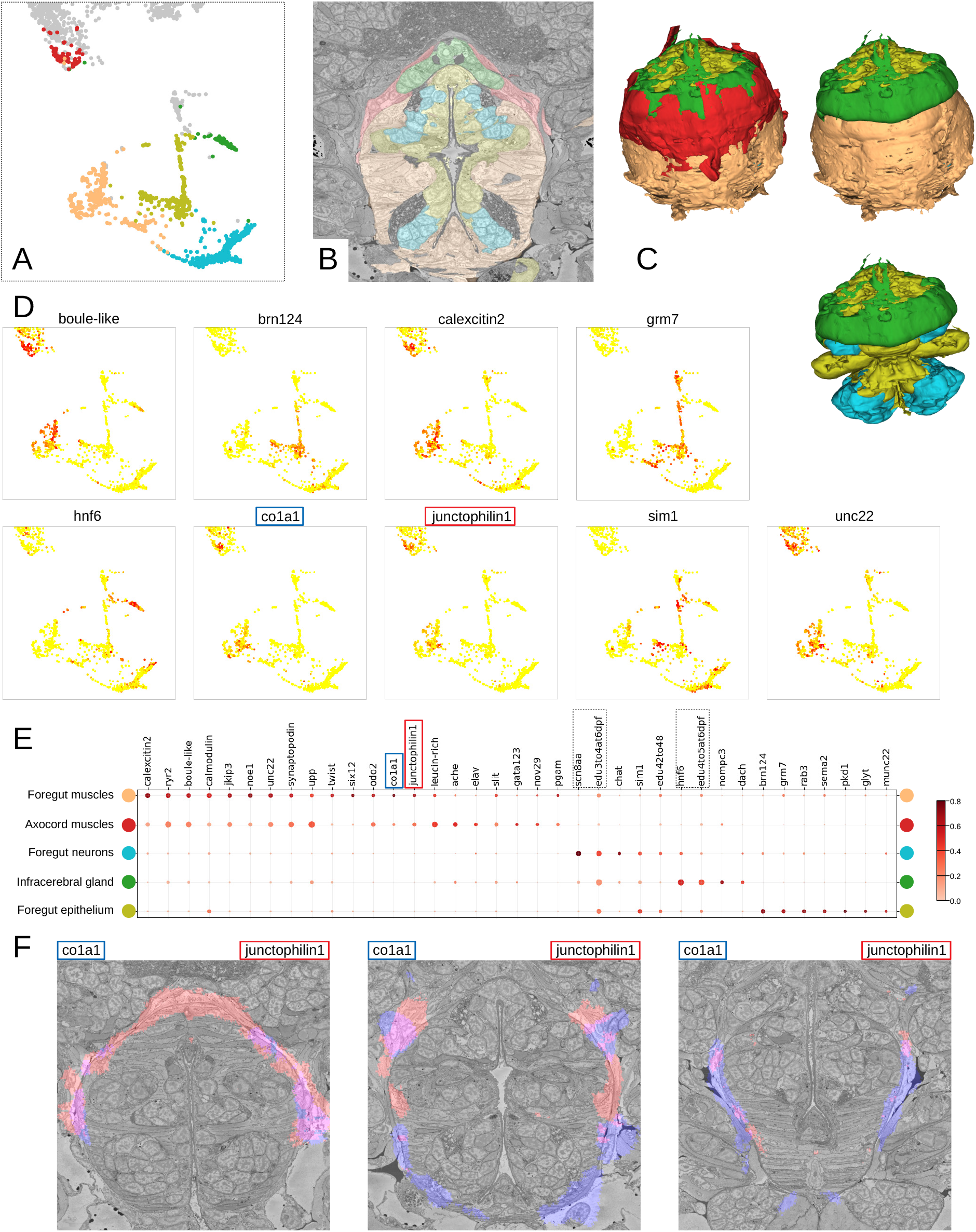
Detailed characterization of the foregut. **A**. Foregut region clusters plotted on the UMAP representation. **B**. The foregut tissues, as defined by the MorphoContextFeatures clustering, shown in the animal volume. **C**. A 3D visualisation of the foregut tissues, as defined by the MorphoContextFeatures clustering: all tissues (upper left panel), surrounding muscles removed (upper right panel), both muscle groups removed (lower panel). **D**. Some of the genes shown to be differentially expressed in the foregut region clusters, plotted on the UMAP representation. **E**. Gene expression dot plot of the foregut region clustering. The size of dots shows how much of the cluster expresses a gene; the colour shows how much of the expression of the gene is confined to a cluster (see Methods). The genes mentioned in the text are enboxed. **F**. The genes specific to the axocord muscles, visualised on the animal volume.

Further inspection revealed that the latter structure is located underneath the brain neuropil and close to the neurosecretory endings of the neurosecretory plexus (***Verasztó et al. (2020)***) and is bounded by thin layers of longitudinal muscle fibres on both the outer and the inner surface (Figure S4A). The location of the structure indicated it might represent the differentiating infracerebral gland - a neurosecretory gland located beneath the brain assumed to play a role in sensing blood glucose levels (***Backfisch (2013)***; ***Baskin (1974)***; ***Hofmann (1976)***; ***Golding (1970)***). The organ is leaf-shaped and lays between the posterior pair of adult eyes (Figure S4B) as also described in (***Backfisch (2013)***) and borders a cavity likely to be a developing blood vessel (Figure S4C). In comparison to the foregut neurons that express sodium channel *scn8aa*, the infracerebral gland cells specifically express homeobox protein *onecut/hnf6* (Figure 9D,E). We also noted that the tissue stains positive for EdU applied between 3 and 5 dpf, indicating that it is still proliferating.

We then identified the prominent muscles surrounding the foregut as the anterior extension of the axochord (***Lauri et al. (2014)***). Visualising the genes specifically expressed in the muscles around the foregut, we noticed specific expression of the type I collagen gene *col1a1*, a marker for axochordal tissue, and of muscle junctional gene *junctophilin1* (Figure 9F). An anterior extension of the axochord around the foregut has been observed in several species and is described in ***Lauri et al. (2014)***; ***Brunet et al. (2016)***; ***Nielsen et al. (2018)***.

## Discussion

We presented an automated method of extracting comprehensive representations of cellular morphology from segmented whole-animal EM data. In stark contrast to the currently available methods, ours refrains both from specifying features explicitly and from predefining a concrete task that might skew the method towards extracting specific features: putting emphasis on some features may bias descriptions towards specific morphologies or cell types, hindering exploratory analysis. To obtain extensive generally useful representations we rely on the latest progress in self-supervised deep learning and set only two criteria for the extracted features: (1) they have to be sufficiently detailed to allow reconstructing the original cell volume and (2) they have to bring visually similar cells in feature space closer to each other than dissimilar ones. We train a deep learning pipeline optimised for these two conditions and show that it produces rich cellular representations, combining shape-, ultrastructure- and texture-based features. More precisely, we show that the obtained representations (MorphoFeatures) capture fine morphological peculiarities and group together cells with similar visual appearance and localization in the animal. Clustering of cells in the MorphoFeature space, and taking advantage of an existing gene expression atlas with cellular resolution, we show that the obtained clusters represent units with differential and coherent gene expression.

Both for morphological type and for tissue detection we illustrate how MorphoFeatures can facilitate unbiased exploratory analysis aimed at detecting and characterising unusual phenotypes or unusual cell neighbourhoods. Moreover, we show quantitative superiority of MorphoFeatures over a broad range of manually defined features on the tasks of cell classification and symmetric partner detection. We also show that, despite being learned implicitly, MorphoFeatures still correspond to visually understandable properties and can often be interpreted. Finally, adding morphological information from the neighbouring cells (MorphoContextFeatures), we reveal the power of our approach to detect local groups of cells with similar morphology that represent tissues.

Since our pipeline is based on the fast-evolving field of self-supervised deep learning, we expect it to further improve with its general advancement. For example, a promising direction are models, such as ***Grill et al. (2020)***, that only rely on positive samples and remove the need to define which cells belong to a different type. Besides MorphoFeatures, we also believe that MorphoContextFeatures, which incorporate neighbourhood information, will potentially benefit from state-of-the-art self-supervised techniques such as contrastive learning techniques on graphs (***Hassani and Khasahmadi (2020)***; ***Velickovic et al. (2019)***). Another interesting direction enabled by the modularity of our pipeline is to integrate data from other modalities. Our features currently integrate shape and texture information of cells, but one could also envision enriching cellular descriptions by adding different types of data, e.g. transcriptomics, proteomics or metabolomics, in a similar manner.

### Versatility of the MorphoFeatures toolbox

We envision our pipeline to enable, for the first time, fast systematic investigation of large EM volumes that involves consistent grouping and characterization of morphological types, discovering particular phenotypes and neighbourhoods, as well as localising cells or tissues in an animal volume that are visually similar to cells or groups of cells of interest. Moreover, we expect MorphoFeatures to be useful for studies where several image volumes of the same species are acquired, be it another sample, a varying phenotype, or a different developmental stage. In such cases, one could envisage using morphological representations to map cells to their corresponding partners in another volume, enabling unbiased comparative analysis at scale. In addition, in case the other volumes are not segmented, one could use the fine texture part of MorphoFeatures to locate patches with phenotypic characteristic signature or cells with distinct ultrastructure (e.g., muscles, secretory cells). Using fine texture descriptors in such cases can also facilitate further segmentation, since the segmentation of the previous volume can be used to define constraints of which textures can co-occur in the cells of the animal. However, given the high sensitivity of neural networks to intensity variations, it will be essential to avoid non-biological systematic change in the visual appearance of cells such as intensity shifts, both within and between the EM volumes. The presented pipeline can also be adjusted for other types of data. For example, since our analysis revealed a high variability in neuronal soma morphologies, it appears reasonable to also apply the pipeline to brain EM volumes, where MorphoFeatures could supplement neuron skeleton features for more precise neuronal morphology description. We also expect the approach to contribute to the analysis of cells in other volumetric modalities, such as X-ray holographic nano-tomography (***Kuan et al. (2020)***). For light microscopy with membrane staining, we expect the shape part of the MorphoFeatures pipeline to be directly applicable and a useful addition to existing analysis tools. Generally, the versatility of the pipeline is enhanced by its modularity that makes it possible to adapt different parts (e.g., cell shape or nucleus fine texture) to datasets where not all the morphological components are desired or can be extracted.

### Identification of cell types and cell type families with MorphoFeatures

Any attempt to define cell types morphologically has so far been hampered by the selection of relevant properties. What are the morphological features that are best suited for recognizing cell types? Regarding neurons, the shapes of dendrites and axons, soma size and spine density have been measured, in combination with physiological properties including resting potential, biophysical properties and firing rate (***Zeng and Sanes (2017)***). Volume EM has recently allowed considerable expansion of morphometric parameters, including various shape, intensity and texture features (***Vergara et al. (2021)***). Using these numerous parameters, it is possible to discern general categories such as neurons, muscle or midgut cells; yet they do not provide further resolution within these groups. Comparing the MorphoFeatures representations to the predefined features (***Vergara et al. (2021)***) hints at supreme resolution power that appears to distinguish morphological groupings down to the cell type level (Figure S3). For example, the subclusters of rhabdomeric photoreceptors of the adult eye and enteric neurons can not be identified in the explicit features representation. Furthermore, the cluster of foregut muscles revealed by the MorphoFeatures appears to be split, with some cells being found closer to striated muscles, and some residing between the groups of neurons and dark neurosecretory cells.

Most important, the cell clusters identified in MorphoFeature space appear to come closest to genetically defined cell types, as demonstrated here, among others, for different types of mechanosensory cells, or cell types of the apical nervous system. Interestingly, the MorphoFeatures also unravel morphological cell types without a genetic match in the cell type atlas (such as the enteric neurons, see above). This is most likely due to the yet limited coverage of the atlas and expected to improve with the continuous addition of genes to the atlas. In particular, the integration of other multimodal omics datasets such as single cell transcriptomics data will be highly rewarding to improve the match between morphological and molecularly defined cell types, which is the ultimate aim of cell type characterization and categorization (***Zeng and Sanes (2017)***).

### Towards an unbiased whole-body identification of tissues and organs

The current revolution in the generation of giant volume EM datasets including representations of entire bodies calls for new strategies in the automated processing and comparison of such datasets. This not only involves segmentation and characterization at the cellular level, but also requires the automated identification of discernable structures at the level of tissues and organs. So far, the recognition of tissues in larger volumes was achieved by manual segmentation. We now present a powerful new algorithm to recognize tissues and organs via their co-clustering in the MorphoContextFeature space, which takes into account similarities between neighbouring cells. Using this algorithm, we reproduce and partially outperform tissue recognition via manual segmentation - as exemplified for the ganglionic nuclei of the nereid brain. Beyond that, we present a comprehensive account of foregut tissues, without prior manual segmentation. This analysis reveals a rich collection of tissues including an anterior extension of the axochord surrounding the foregut (***Lauri et al. (2014)***) and the infracerebral gland (***Baskin (1974)***; ***Golding (1970)***), an adenohypophysis-like neurosecretory tissue that we identify as anterior bilateral extensions of the foregut roof.

While the manual segmentation of tissues mostly relies on the detection of tissue boundaries, our automated tissue detection equally processes the morphological similarities of its constituting cells. This enables tissue separation even in cases where morphological boundaries appear to be absent, as is the case for the anterior and posterior dorsal ganglionic nuclei, which are lacking a clear boundary but are composed of morphologically different cell types that are also genetically distinct (***Vergara et al. (2021)***).

## Conclusion

The revolution in multi-omics techniques and volume electron microscopy has unleashed enormous potential in generating multimodal atlases for tissues as well as entire organs and animals. The main unit of reference in these atlases is the cell type, which has been defined, and is manifest, both molecularly and morphologically. Our work now provides a versatile new tool to recognize cell types in an automated, unbiased and comprehensive manner solely based on ultrastructure and shape. For the first time, the MorphoFeatures will allow us to compare and identify corresponding cell types in datasets of the same species, but also between species, on purely morphological grounds.

Furthermore, the MorphoContextFeatures automatically recognize tissues and organs across entire bodies, opening up new perspectives in comparative anatomy. Without previous knowledge, we can obtain the full complement of tissues and organs for a given specimen, and quantitatively compare them to those of other specimens of the same or other species. In combination with cell type-, tissue-, and organ-specific expression data this will enable us to identify corresponding traits in an unbiased manner and with unprecedented accuracy.

## Methods and Materials

### Data and preprocessing

#### Data

We used the dataset of the marine annelid Platynereis dumerilii at late nectochaete stage (6 dpf) from ***Vergara et al. (2021)***. The dataset comprised an SBEM volume of the complete worm in 10 × 10 × 25 nm resolution, 3D cell and nucleus segmentation masks as well as WMISH gene expression maps. The segmentation contains 11,402 cells with nuclei that belong to multiple genetically and morphologically distinct cell types. It is important to note that for neurons only somata were segmented, since the imaging resolution did not allow for neurite tracing. The gene expression maps cover mainly differentiation genes and transcription factors, which were registered onto the EM volume, exploiting high stereotypy of the platynereis at this stage of development. Gene expression values were also assigned to the EM segmented cells. The maps contain 201 genes and 4 EdU proliferation stainings. We refer to ***Vergara et al. (2021)*** for further details. Since the intensity correction algorithm that was used in the EM volume performed unreliably in regions close to the boundaries of the volume in the z-axis, we excluded all cells located within the region of uncertain intensity assignment quality. These regions correspond to parts of the animal’s head and the pygidium.

#### Preprocessing

The EM dataset is used to extract the three morphological components separately for cytoplasm and nuclei as follows. Due to the limitations of the GPU memory, for the coarse texture a central crop is taken from a downsampled version of the data (80 × 80 × 100 nm). The centre of mass of the nucleus is determined, and a bounding box of size 144 × 144 × 144 pixels (cytoplasm) or 104 × 104 × 104 (nuclei) is taken around it. During prediction the bounding box is increased to 320 × 320 × 320 (cytoplasm) and 160 × 160 × 160 (nuclei) to fit more information. Fine texture patches have a higher resolution (20 × 20 × 25 nm) and are taken separately from cytoplasm and nucleus.

For this the bounding box surrounding cytoplasm or nucleus is split into cubes of 32 × 32 × 32 pixels, and only those cubes are considered, which are less than 50% empty. For both coarse and fine texture the data is normalised to the range of [0, 1] and for cytoplasm crops the nuclei are masked out and vice versa.

In order to obtain point clouds and normals of the membrane surfaces, we first converted the segmented 3D voxel volumes into triangular meshes. More specifically, we first applied the marching cubes algorithm (***Lorensen and Cline, 1987***; ***Van der Walt et al., 2014***) followed by Laplacian smoothing (***Schroeder et al., 1998***) to obtain smooth triangular surface meshes. These meshes were then simplified and made watertight using ***Stutz and Geiger (2020)***; ***Cignoni et al. (2008)***. Having obtained smooth triangular meshes enables us to compute consistent normals and to apply complex deformations to the underlying surface. During training and inference, the meshes and their surface normals are sampled uniformly to be then processed by the neural network.

### Unsupervised deep learning pipeline

This section describes our proposed algorithm in more detail. On a high level, our model consists of six encoder networks which leverage graph convolutional as well as regular convolutional layers in order to learn shape and texture aspects of a cell’s morphology. We are using self-supervised learning in combination with stochastic gradient descent in order to obtain good network parameters. The objective that is optimised is a weighted sum of the NT-Xent loss (***Chen et al. (2020)***) and the mean squared error and can be interpreted as an unbiased proxy task. We will detail and elaborate on these aspects in the following sections.

#### Neural network model

The proposed network, as depicted in (Figure 1A), contains six encoders which are encoding shape, coarse and fine-grained texture of cells and nuclei.

The shape encoder takes as input a set of points in the form of 3D Euclidean coordinates as well as the corresponding normals (see Data and preprocessing) and outputs a N-dimensional shape feature vector. The network itself is a DeepGCN model (***Li et al. (2019)***), which is a deep neural network for processing unstructured sets such as point clouds. Conceptually it consists of three building blocks. In the first block, k-nearest neighbour graphs are built dynamically in order to apply graph convolutions. This can be interpreted as pointwise passing and local aggregation of geometric information which is repeated multiple times. In the second block, the pointwise aggregated geometric information is fused by passing it through a convolutional layer followed by a max pooling operation that condenses the information into a high-dimensional latent vector. Finally, the obtained latent vector is passed through a multilayer perceptron to obtain the shape component of the overall feature vector.

The encoding of fine texture details and coarser contextual information is done using the encoder of a U-Net (***Ronneberger et al. (2015)***). The fully convolutional network reduces spatial information gradually by employing 3 blocks that consist of 3D convolutional layers with ELU activations (***Clevert et al. (2015)***) separated by 3DMaxPooling layers. The number of feature maps starts with 64, increases twofold in each block, and afterwards gets reduced to 80 by an additional convolutional layer followed by an ELU activation. In the end a global average pooling operation is applied to reduce feature dimensionality from 3D to 2D. To ensure that empty regions of the data are not affecting the final feature vector, they are excluded for this final pooling.

#### Training

In order to find suitable representations we leverage self-supervised contrastive learning (CL) (***Chen et al., 2020***; ***Hadsell et al., 2006***). The idea is to create a representation space in which morphologically similar cells are embedded close to each other, while dissimilar ones are embedded far apart. For learning representations of shape, we follow a purely contrastive approach, whereas for texture representations we combine the CL objective with an Autoencoder reconstruction loss.

#### Both are described in more detail below

The training procedure for CL is the same for shape and texture representations. First, we randomly sample a mini-batch of cells or nuclei. Two augmented views are then created for each sample in the batch (see Data augmentations) and concatenated to form an augmented batch of twice the size. The augmented batch is passed through the neural network encoder to compute representations. Those representations that share the same underlying sample are considered as positive pairs whereas the others are considered as negative pairs. The NT-Xent loss (***Chen et al., 2020***) is then used to score the representations based on their similarity, and, therefore, encourages the network to embed positive pairs closer to each other than negative pairs.

For learning shape representations, we first augmented all cell and nucleus meshes using biharmonic deformation fields and as-rigid-as-possible transformations in a preprocessing step (see Data augmentation). More specifically, we computed 10 random views for each membrane surface. The augmented meshes are then used for training, i.e. we compute surface normals and sample 1024 points and their corresponding normals which form the inputs of the shape encoders. In each training iteration, we sampled 48 cells/nuclei, which resulted in a batch size of 96. Adam (***Kingma and Ba, 2014***) with a learning rate of 0.0002 and a weight decay of 0.0004 was used to update the network parameters. The hyperparameters were obtained by a random search.

In order to learn representations of texture, we combine the NT-Xent loss with an Autoencoder reconstruction objective. The latent representations, taken before the dimensionality reduction step of global average pooling, are further processed by a symmetric decoder part of the UNet network with only 1 block, which aims to reconstruct the network inputs. The reconstructions are then compared to the original inputs and scored by a mean squared error loss. Additionally, an L2-Norm loss was applied to flattened bottleneck features to restrict the range of possible values. The final loss is thus a weighted sum of the NT-Xent, the mean squared error loss and the L2-Norm loss. The batch size was set to 12 and 16 for cell and nuclei coarse texture and to 32 for fine texture. For training we used Adam (***Kingma and Ba, 2014***) as optimizer with a learning rate of 0.0001 and a weight decay of 0.00005. The model was evaluated every 100 iterations and the best result was saved with a patience of 0.95. The learning rate was reduced by 0.98 whenever the validation score did not show improvement with a patience of 0.95.

#### Data augmentation

A key component for successful contrastive representation learning are suitable data augmentations. In our context, applying augmentations to a cell can be interpreted as creating morphological variations of a cell. By creating positive and negative pairs (cf. previous section), the network learns to distinguish true variations from noise and non-essential ones. It is therefore important to find a suitable set of transformations that distort the right features of a cell and do not change its morphological identity.

As augmentations for shape we leveraged biharmonic and as-rigid-as-possible transformations (***Botsch and Kobbelt, 2004***; ***Sorkine and Alexa, 2007***) as they deform cellular and nuclear membranes in a way that retains most characteristic geometric features of the original surface. In order to compute these deformations, we sample so-called handle regions on the membrane surface. These regions are translated in a random, normal or negative normal direction. The deformation constraint is then propagated to the rest of the surface either via a biharmonic deformation field or an as-rigid-as-possible deformation. We combine these deformations with random anisotropic scaling, rotations and/or symmetry transformations.

To create positive samples for the contrastive training of texture we used random flips and rotations and elastic transformations of the data.

#### Inference

In order to compute the MorphoFeatures representation for the whole animal, we processed each cell’s cellular and nuclear shape and texture, as described in previous sections, with our trained network. Each cell is represented by a 480-dimensional vector that consists of a nuclear and cellular part. Each of these parts consists of a shape, coarse and fine-grained texture 80-dimensional feature vector.

### Quantitative feature evaluation

In this section we describe our feature evaluation experiments in more detail. In order to assess the quality of MorphoFeatures quantitatively we conducted cell-type prediction and bilateral pair analysis experiments. We use the same set of explicit features that were used in ***Vergara et al. (2021)*** for analysing cellular morphology as a baseline for comparisons.

#### Explicit feature baseline

The explicitly defined representation includes shape features (volume in microns, extent, equivariant diameter, major & minor axes, surface area, sphericity, max radius), intensity features (different quantiles of intensity mean, median and standard deviation) and Haralick texture features. These features were calculated, whenever appropriate, for cell cytoplasm, nucleus and chromatin segmentation. The calculated features were taken from https://github.com/mobie/platybrowser-project/blob/main/data/1.0.1/tables/sbem-6dpf-1-whole-segmented-cells/morphology.tsv for cellular segmentation and from https://github.com/mobie/platybrowser-project/blob/main/data/1.0.1/tables/sbem-6dpf-1-whole-segmented-nuclei/morphology.tsv for nucleus and chromatin segmentation. The UMAP of these features was taken from https://github.com/mobie/platybrowser-project/blob/main/data/1.0.1/tables/sbem-6dpf-1-whole-segmented-cells/morphology_umap.tsv.

#### Classification of visibly distinct groups of cells

In order to verify that MorphoFeatures can distinguish visibly different groups of cells, we evaluated the prediction accuracy of a logistic regression model that was trained on a small dataset of 390 annotated cells taken from ***Vergara et al. (2021)*** (Figure 2A). The original annotation contained 571 neurons, 141 epithelial cells, 55 midgut cells, 53 muscle cells, 51 secretory cells, 41 dark cells and 25 ciliated cells. We first corrected some cells with clearly wrong assignments. Then for the classes that contained less than 60 cells we manually annotated more cells. From bigger classes we selected 60 cells, selecting the most morphologically diverse ones when possible. For ciliated cells we were only able to locate 30 cells in the whole animal.

First we computed the MorphoFeatures representation for each cell in the animal (excluding cells with severe segmentation errors) and standardised features across the whole dataset. We then trained and validated a logistic regression model using a stratified k-folds cross validator with 5 folds. We used the implementations of logistic regression, stratified k-fold from Scikit-learn (***Pedregosa et al., 2011***) with the following arguments: C=1, solver=‘lbfgs’. To prevent potential overfitting due to the high number of features we used feature agglomeration (***Pedregosa et al., 2011***) (n_clusters=100) to reduce the amount of similar features.

#### Bilateral pair analysis

The criterion of classifying distinct groups of cells in the previous section can be seen as a coarse and global way of evaluating MorphoFeatures. In this section we describe how we used the animal’s bilateral symmetry to have a more fine-grained measure for quantitatively evaluating nuanced morphological differences. The measure, which we will refer to as bilateral distance, was introduced in ***Vergara et al. (2021)*** and leverages the fact that most cells in the animal have a symmetric partner cell on the other side of the mediolateral axis. The bilateral distance is computed in the following way: first, we determine for each cell *c* a group of N potential partner cells {*c*_1_, …, *c*_*N*_} on the opposite side. We then compute a ranking based on the Euclidean distances ‖*c* − *c*^′^‖ in feature space between cell *c* and all other cells *c*^′^ ≠ *c*. Since *c*’s symmetric partner has the same morphology, up to natural variability, it should be among the closest cells in the ranking. We then determine the highest ranked partner cell as rank(‖*c* − *c*_*n*_‖). In doing this for all cells, we create a list containing the ranking positions of cells to their symmetric partner. The bilateral distance is then defined as the median of this list and can be interpreted as a measure of how close two morphologically very similar cells are on average in feature space.

### Adding neighbour morphological information

This section describes how neighbourhood information was incorporated into MorphFeatures in order to obtain MorphoContextFeatures. First we built a region adjacency graph from the cell segmentation masks using scikit-image (***Van der Walt et al., 2014***). The obtained graph reflects the animal’s cellular connectivity, i.e. cells are represented by nodes in the graph and nodes are connected by an edge if the corresponding cells are neighbours. We then assign node attributes by mapping the MorphoFeatures representation of a cell to the corresponding node. Neighbourhood features are then obtained by computing the mean over a node’s MorphoFeatures vector and the MorphoFeatures vectors of its surrounding neighbours. Finally, a cell’s neighbourhood features are concatenated with the cell’s MorphoFeatures to obtain the MorphoContextFeatures representation.

### Feature analysis

#### Visualising features

In order to visualise the features extracted by our pipeline (Figure 3) we manually selected a set of features that visually corresponded to potentially interpretable textures/shapes and showed their extremes with cells that have the respective minimal/maximal value of this feature. The cells with obvious segmentation errors that resulted in incorrect shapes or textures were ignored. In order to visualise the 3D cells and nuclei we selected a 2D view that we presume shows the shape or texture feature common between all 4 selected cells with minimal or maximal value of the feature.

#### UMAP representation analysis

Visualisation was done using the UMAP package (***McInnes et al. (2018)***) with the following parameters: euclidean metric, 15 neighbours and 0 minimal distance. Cell metadata, such as annotated types, animal regions or gene expression were plotted on the resulting UMAP.

#### Clustering

Cell clustering was done according to the following procedure. First the features were standardised to zero mean and unit variance. Then, following (***Dubourg-Felonneau et al., 2021***) a weighted k-neighbour graph was constructed from MorphoFeatures representations of all the animal cells using the UMAP algorithm (***McInnes et al., 2018***) with the following arguments: n_neighbors=20, metric=‘euclidean’, min_dist=0. Afterwards the Leiden algorithm (***Traag et al., 2019***) for community detection was used to partition the resulting graph, using the CPMVertexPartition method and a resolution parameter of 0.004. This resulted in 16 clusters, one of which was discarded, since it contained only cells with split segmentation errors. For more fine-grained clustering the resolution was increased to 0.1 to split apart secretory and ciliary band cells, 0.2 to split photoreceptors and enteric neurons, 0.07 to split the midgut cluster, 0.005 for muscle neighbourhood cluster (n_neighbors for the UMAP clustering was adjusted to 10) and 0.01 for splitting foregut neurons and epithelial cells. To visualise average shape and texture of a cluster (5B,C), a median feature vector was computed across the cluster and the cell with the feature vector closest to the median one was taken as the average cell.

#### Gene expression analysis

To produce the cluster gene expression ‘dot plots’ (Figure 5A, 6B, 8C, 9E) the gene expression values for each cell were taken from ***Vergara et al. (2021)*** at https://github.com/mobie/platybrowser-project/blob/main/data/1.0.1/tables/sbem-6dpf-1-whole-segmented-cells/genes.tsv. Similar to ***Vergara et al. (2021)***, for each cluster and for each gene we calculated three values: (A) the mean expression in the cluster, (B) the fraction of the total animal/region expression within this cluster (the sum of expression in this cluster divided by the sum of the expression in the whole animal or the selected region) and (C) the specificity defined as *C* = 2*AB/*(*A* + *B*). Intuitively, *A* indicates how much of the cells in a given cluster express a given gene, *B* shows how selective this gene for this cluster is and *C* is a harmonic mean of the two values. To create a dot plot only the genes with *C* value above 0.15 were used. For the neuron clusters 1, 3 and 5 (Figure 5A) the threshold was increased to 0.2 due to a high number of specifically expressed genes available. For the midgut clusters (Figure 6) to show only the genes differing between the two clusters, the genes were removed, where the specificity was above 0.15 in both clusters, but differed by less than 0.4. The size of dots in the plots reflects *A* and the colour corresponds to *B*. The cluster with the most gene expression was determined and the remaining clusters were sorted by their similarity to it. To illustrate some of the genes showing differential expression in these groups (Figure 5, 8, 9), the corresponding regions of the UMAP representation were cut out and these genes were plotted on top.

#### Ganglionic nuclei

To compare ganglionic nuclei (Figure 8) we used manually segmented regions from https://github.com/mobie/platybrowser-project/blob/main/data/1.0.1/tables/sbem-6dpf-1-whole-segmented-cells/ganglia_ids.tsv and https://github.com/mobie/platybrowser-project/blob/main/data/1.0.1/tables/sbem-6dpf-1-whole-segmented-cells/regions.tsv and gene clusters from https://github.com/mobie/platybrowser-project/blob/main/data/1.0.1/tables/sbem-6dpf-1-whole-segmented-cells/gene_clusters.tsv (***Vergara et al. (2021)***).

### Data visualisation

We used matplotlib (***Hunter, 2007***) and seaborn (***Waskom, 2021***) for plotting the data, MoBIE (***Vergara et al., 2021***) for visualising cells in the EM volume and vedo (***Musy et al., 2022***) for visualising meshed cells and tissues.

## Acknowledgments

We thank all the members of the Kreshuk and Uhlmann group, especially Constantin Pape, for helpful comments. We further thank Christian Tischer for implementing features essential for biological analysis and exploration in MoBIE. We thank Hernando M. Vergara for assistance in cell type recognition and Kimberly I. Meechan for help with comparison to explicit features. We would also like to acknowledge the support of the EMBL IT Services, especially Juri Pecar for maintaining the GPU resources. The work was supported by EMBL internal funds, and by a grant from the European Research Council (NeuralCellTypeEvo 788921) to D.A.

## Supplementary

**Figure S1.**
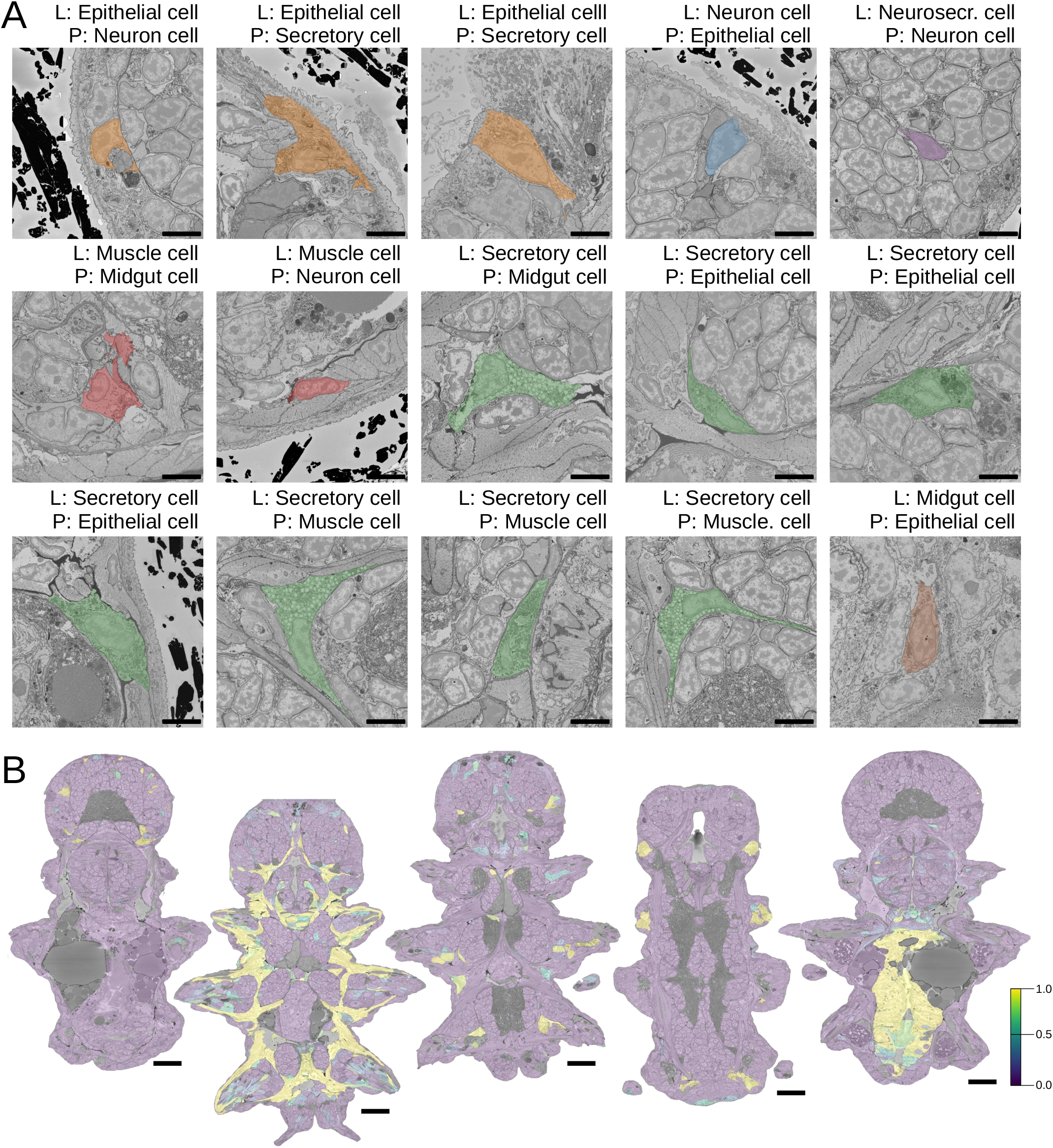
Morphological class predictions. **A**. All the errors made during any of the cross-validation runs of the logistic regression model (L – True Label, P – Prediction, scale bars: 5 *µ*m). **B**. Whole-animal prediction of (from left to right) dark neurosecretory, muscle, secretory, ciliary band and midgut cells (scale bars: 25 *µ*m).

**Figure S2.**
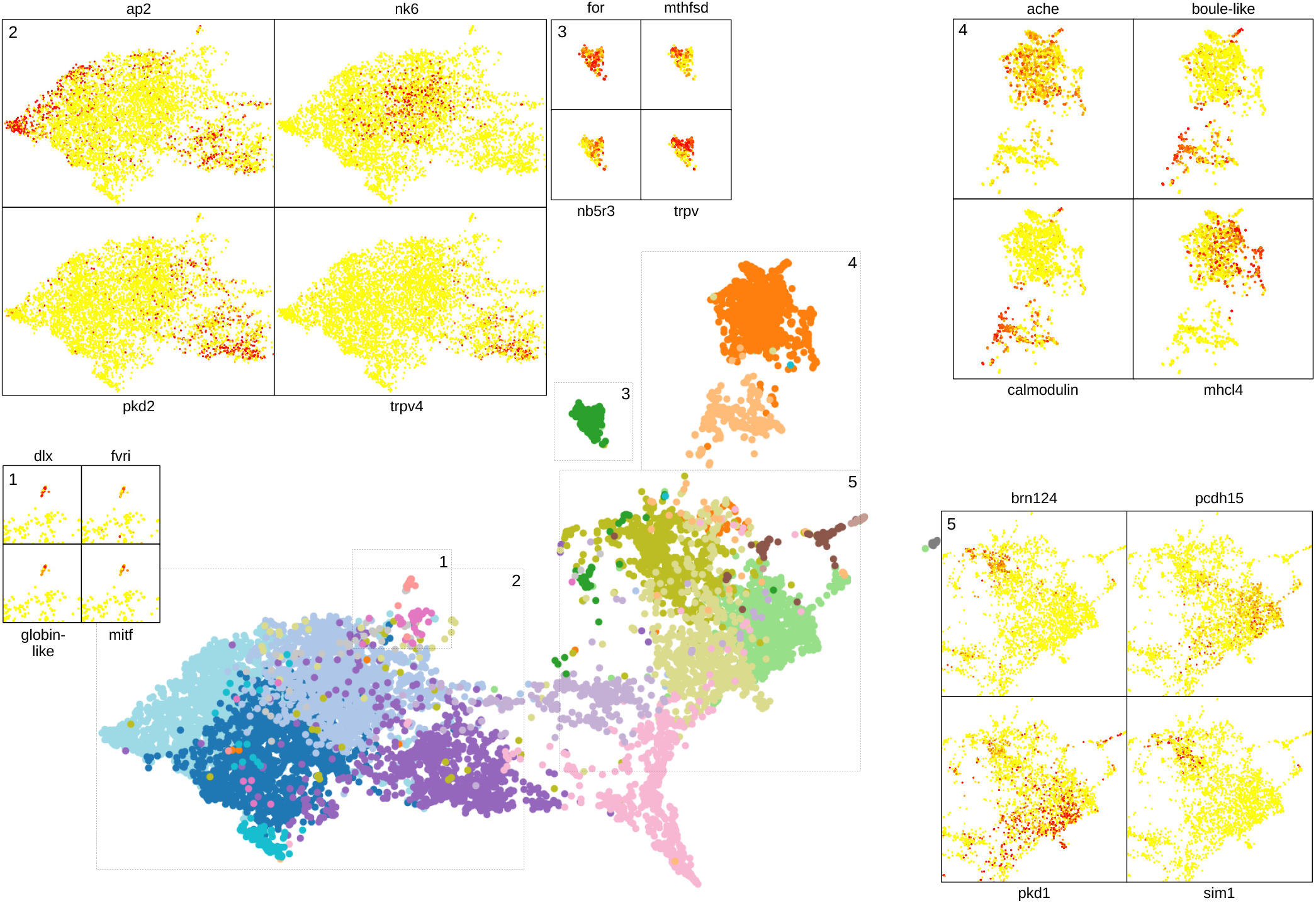
Clustering and gene analysis. Some of the genes shown to be differentially expressed in neuron, midgut, muscle and epithelial clusters and in the photoreceptor cells are visualised on the UMAP representation.

**Figure S3.**
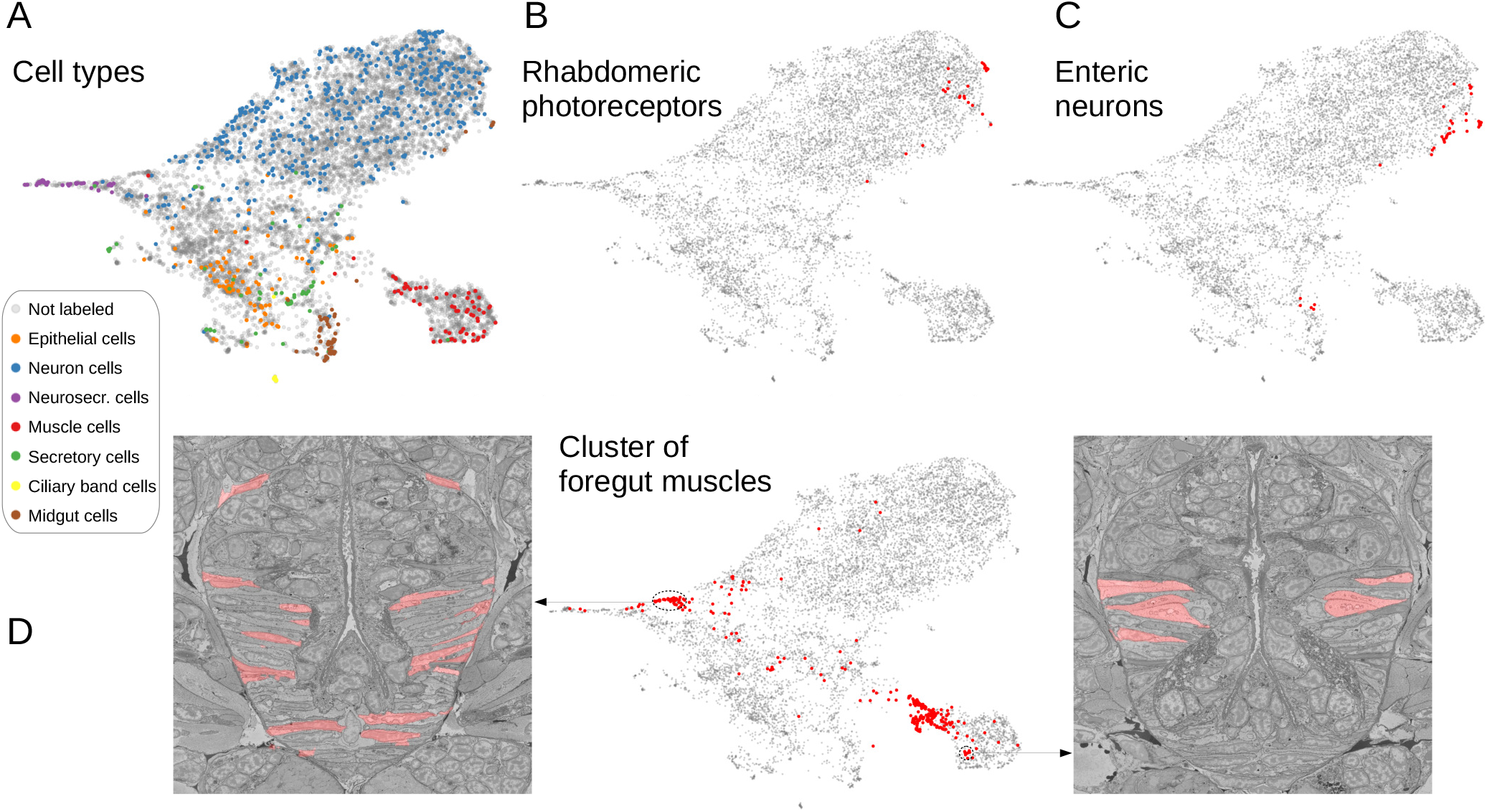
Comparing MorphoFeatures to a set of manually defined features from ***Vergara et al. (2021)***. **A**. The set of manually defined features is visualized in 2D using UMAP. The cells for which annotations are available are visualized in respective colors. **BC**. The MorphoFeatures cluster of (**B**) the rhabdomeric photoreceptors and (**C**) enteric neurons is visualised on the UMAP representation of the manually defined features, showing a bigger spread of cells across the morphological space. **D**. The MorphoFeatures cluster of the foregut muscles is split into multiple groups on the UMAP representation of the manually defined features. For two groups, cells are visualised in the animal volume, showing similarity of the cells comprising them.

**Figure S4.**
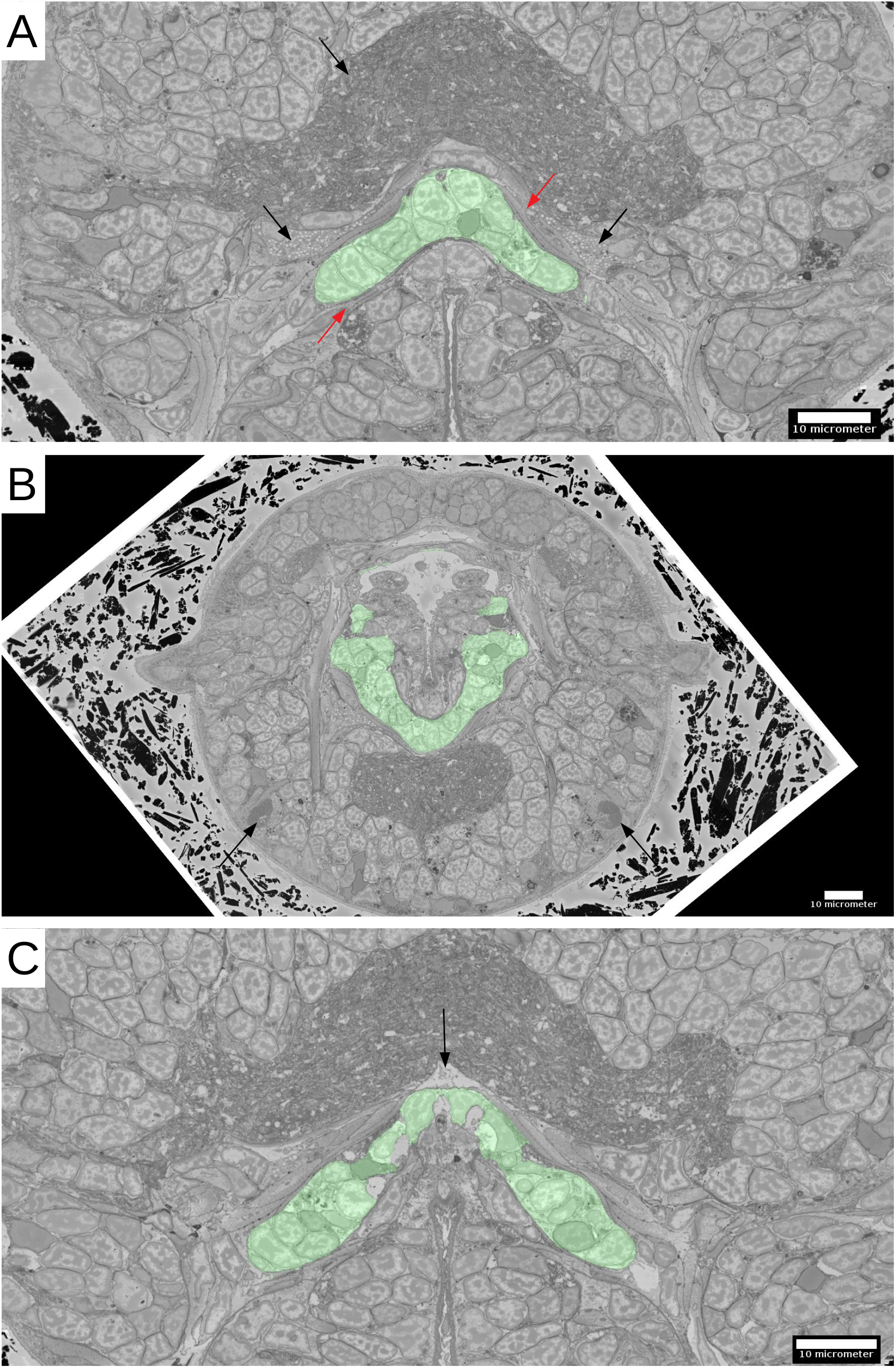
Infracerebral gland. **A**. The location of the gland in the head. The neuropil and the secretory cells are pointed up by black arrows, the surrounding muscle layers - by red arrows. **B**. The shape of the gland and its position relative to the posterior pair of adult eyes (black arrows). **C**. A cavity likely to be a developing blood vessel (black arrow) on top of the gland.

